# Generation and characterization of a barley strigolactone mutant collection: from plant architecture to drought stress response

**DOI:** 10.64898/2026.05.28.728395

**Authors:** Irene M. Fontana, Weronika Buchcik, Jochen Kumlehn, Michael Melzer, Goetz Hensel, Agata Daszkowska-Golec, Marek Marzec

## Abstract

Strigolactones (SLs) are known to regulate shoot architecture and to be involved in plant responses to environmental stress, whereas their specific contributions to drought adaptation in barley remain incompletely defined. In this study, we analysed transcriptional, hormonal, and physiological responses to water deficit in barley SL mutants affected in early biosynthesis (*Hvd10* and *Hvd17*), late biosynthesis (*Hvmax1a*), or signalling (*Hvd14*). The *Hvd10, Hvd17*, and *Hvd14* mutants exhibited the typical high-tillering phenotype of SL deficiency, whereas *Hvmax1a* displayed characteristics similar to the wild type (WT), indicating functional differences within the SL biosynthetic pathway.

Transcriptome analysis showed a clear overlap in gene expression among the high-tillering SL mutants under both control and drought conditions. We also used computational methods to identify potential transcription factors that might regulate SL-dependent gene expression. A drought experiment showed that SL mutants exhibited reduced biomass, relative water content, and photosynthetic efficiency, with the most pronounced effects observed in the high-tillering lines. Drought also activated the abscisic acid (ABA) pathway in all genotypes, with particularly high accumulation of ABA metabolites in the high-tillering SL mutants. Notably, *Hvmax1a* resembled the mutant-like metabolic profile, despite maintaining a wild-type-like architecture.

Taken together, these results provide new insights into the roles of SL pathway components in drought responses and highlight functional differences among individual genes influencing both plant architecture and stress-related transcriptional programmes. Furthermore, the mutants generated in this study using Cas9-mediated genome editing represent a valuable genetic collection for future research into SL-mediated development and stress responses in barley.

## Introduction

Strigolactones (SLs) are a group of carotenoid-derived phytohormones characterized by the presence of a butenolide moiety (Al-Babili and Bouwmeester, 2015). They were initially identified as signalling compounds that induce seed germination in root-parasitic plants, and were subsequently found to stimulate hyphal branching in arbuscular mycorrhizal (AM) fungi (Akiyama *et al*., 2005; Cook *et al*., 1966). Beyond their role in rhizosphere signalling, SLs inhibit shoot branching in both monocotyledonous and dicotyledonous plants (Gomez-Roldan *et al*., 2008; Umehara *et al*., 2008). They also regulate various developmental processes, including root system architecture, leaf senescence(Koltai, 2011; Yamada *et al*., 2014), and plant responses to environmental (Marzec *et al*., 2013; Marzec and Muszynska, 2015; Yoneyama, 2019).

To date, more than 35 structurally distinct SLs have been identified across various plant taxa (Seto, 2023). The first SL to be isolated, strigol, was found to have a characteristic four-ring skeleton (Cook *et al*., 1966). Compounds sharing this structural feature are collectively called canonical SLs (Waters *et al*., 2017; Yoneyama and Brewer, 2021). These molecules feature a tricyclic lactone moiety (ABC rings, arranged in a 6-5-5 or 7-5-5 configuration) connected via an enol ether bridge to a butenolide (D ring), which invariably adopts the 2′R configuration in natural SLs (Mashiguchi *et al*., 2021). Based on the stereochemistry at the BC-ring junction, canonical SLs are further classified into two groups: orobanchol-type (α-orientation) and strigol-type (β-orientation) (Xie *et al*., 2013). Recently, several SLs lacking the typical ABC ring structure have been identified and are referred to as non-canonical SLs (Yoneyama *et al*., 2018). Evidence now suggests that canonical SLs primarily act as rhizospheric signals, whereas non-canonical SLs are considered the main hormonal regulators of plant architecture (Wang *et al*., 2024). Furthermore, these two groups are believed to originate from divergent biosynthetic pathways that emerged during evolution (Zhou *et al*., 2025).

SLs originate from the carotenoid pathway, where all-trans-β-carotene is isomerised and cleaved through the combined action of DWARF27 (D27), CAROTENOID CLEAVAGE DIOXYGENASE 7 (CCD7), and CAROTENOID CLEAVAGE DIOXYGENASE 8 (CCD8) to produce the central bicyclic precursor carlactone (CL) (Alder *et al*., 2012; Lin *et al*., 2009; Seto *et al*., 2014; Waters *et al*., 2012). CL is then oxidised to carlactonoic acid (CLA) by a cytochrome P450 of the CYP711A family encoded by *MORE AXILLARY GROWTH1* (*MAX1*) (Abe *et al*., 2014; Seto *et al*., 2014). This early and fundamental pathway, shared by all plants, supplies the universal intermediates CL and CLA, which serve as branching points for the diversification of SL structures (Yoneyama and Brewer, 2021). Beyond CLA, downstream modifications mediated by CYP711A homologs and other P450 enzymes vary across species, leading to structurally and functionally diverse SLs (Kodama *et al*., 2022; Wakabayashi *et al*., 2019).

Core SL biosynthesis mutants show a full spectrum of SL-related traits, including excessive tillering, dwarfism, and decreased root exudation (Chen *et al*., 2023; Ito *et al*., 2022). In contrast, several downstream mutants exhibit impaired production of specific SLs but do not display the branching phenotype typical of core pathway mutants (Wakabayashi *et al*., 2019; Zhou *et al*., 2025). These results suggest that the conserved core pathway regulates shoot branching, while species-specific downstream pathways branch out to expand SL diversity. Although the core SL biosynthetic pathway is now well understood, the diversification of SLs remains unclear, with new enzymes continually being identified (Zhou *et al*., 2025).

A similar situation applies to SL signalling, where the early steps of signal transduction are relatively well understood, whereas our knowledge of downstream components, such as SL-dependent transcription factors, remains limited (Korek and Marzec, 2024). The SL signalling pathway operates through a ubiquitin-dependent degradation system. The receptor DWARF14 (D14), an α/β-hydrolase, perceives SLs and initiates degradation of repressor proteins, including DWARF53 (D53) in *Oryza sativa* (rice) and SUPPRESSOR OF MAX2 1-LIKE6/7/8 (SMXL6/7/8) in *Arabidopsis thaliana* (Arabidopsis), in a process that requires the F-box protein DWARF3 (D3)/MORE AXILLARY BRANCHES 2 (MAX2) (Arite *et al*., 2009; Soundappan *et al*., 2015; Wang *et al*., 2015; Zhou *et al*., 2013). D14 is unusual among plant hormone receptors in that it possesses catalytic activity toward its ligand, adding complexity to the signalling mechanism and raising debate over whether hydrolysis is required for signal (Hamiaux *et al*., 2012; Seto *et al*., 2019; Yao *et al*., 2016). Structural studies suggest that ligand binding induces conformational changes in D14 that promote complex formation with signalling partners, although alternative models propose that the intact SL molecule itself drives this process (Shabek *et al*., 2018; Yao *et al*., 2016). Despite significant progress, the exact relationship between SL hydrolysis, receptor conformational dynamics, and signal initiation remains unclear (Marzec and Brewer, 2019).

Genetic mutants have played a crucial role in advancing our understanding of SL biology. Early studies in Arabidopsis, pea, rice, and petunia showed that excessive branching phenotypes resulted from defects in SL biosynthesis or signalling, long before SLs were identified as plant hormones (Beveridge and Kyozuka, 2010; Dun *et al*., 2009; Matusova *et al*., 2005). In barley, the creation and analysis of SL-related mutants have similarly provided important insights into SL function. For example, mutations in *HvD53A*, a negative regulator of SL signalling, impair photosynthesis but enhance drought tolerance, highlighting the complex role of SLs in coordinating growth and stress responses (Korek, Buchcik, *et al*., 2025). Similarly, studies of mutants in LATERAL BRANCHING OXIDOREDUCTASE (LBO), an enzyme involved in SL biosynthesis, have shown how specific biosynthetic steps regulate shoot branching (Inoue *et al*., 2025). Transcriptomic and multi-omic studies of the SL-insensitive receptor mutant *Hvd14.d* (Marzec *et al*., 2016) have further shown wide-ranging effects on hormonal balance, stress adaptation (Daszkowska-Golec *et al*., 2023; Korek, Mehta, *et al*., 2025), and yield (Kelly *et al*., 2025), supporting the diverse functions of SLs. The goal of this study was to create a collection of barley mutants using Cas endonuclease-mediated genome editing, specifically addressing key components of the SL biosynthesis and signalling pathways, thus offering a genetic toolset to explore the roles of these hormones in plant development and stress responses. We exemplify the usefulness of this mutant collection by investigating how SLs influence drought responses. Since barley is both a model cereal and a major crop, establishing a comprehensive collection of SL mutants is of particular importance, laying the foundation for breeding approaches aimed at improving yield stability and stress resilience.

## Results

### Cas endonuclease-mediated generation of barley mutants in strigolactone biosynthesis and signalling

Loss-of-function alleles in specific SL pathway genes were generated in barley using RNA-guided endonucleases. Guide RNAs were designed to target the early coding regions of each gene, avoiding areas with alternative in-frame start codons to ensure effective disruption of translation. Sequencing of regenerated plants revealed multiple insertions and deletions at the expected target sites, mainly causing frameshift mutations. Most mutations were stably inherited, and homozygous individuals were identified through genotyping. For each gene, four independent homozygous allelic lines were chosen for further analysis. Segregation of the editing construct was monitored in subsequent generations, and *cas9* and gRNA transgene-free lines were identified. All lines examined in this study belonged to the M3-M6 generations.

For *HvD17*, editing of four target motifs resulted in allelic variants carrying frameshift mutations. Three lines proved transgene-free, while one retained the editing construct.

For *HvD10*, mutagenesis at two selected target motifs produced alleles predicted to impair protein function. Of the four lines, three are transgene-free, while one still contains the editing construct.

For *HvMAX1a*, addressing four target motifs succeeded in loss-of-function alleles. All lines were verified to be transgene-free.

For *HvD14*, targeted editing of four specific motifs resulted in four independent allelic variants with frameshift mutations. The absence of the editing construct was confirmed in all lines.

The mutations identified in the generated mutant lines are summarised in Supplementary Table S1, and the positions of the Cas9 target motifs within the addressed genes are shown in Supplementary Fig. S1).

### Phenotypic consequences of mutations in strigolactone biosynthesis and signalling genes in barley

We initially compared plant architecture between WT and the analysed mutant lines grown under control conditions. Clear genotype-dependent differences were identified in both plant height and shoot branching. WT plants had an average height of 87.5 cm, while *Hvd17*, *Hvd10*, and *Hvd14* mutants were significantly shorter, with mean heights of 67.7, 66.4, and 65.1 cm, respectively, representing 77%, 76%, and 74% of WT. In contrast, *Hvmax1a* plants remained similar to WT, with a mean height of 87.2 cm (Fig. 1).

**Figure 1.**
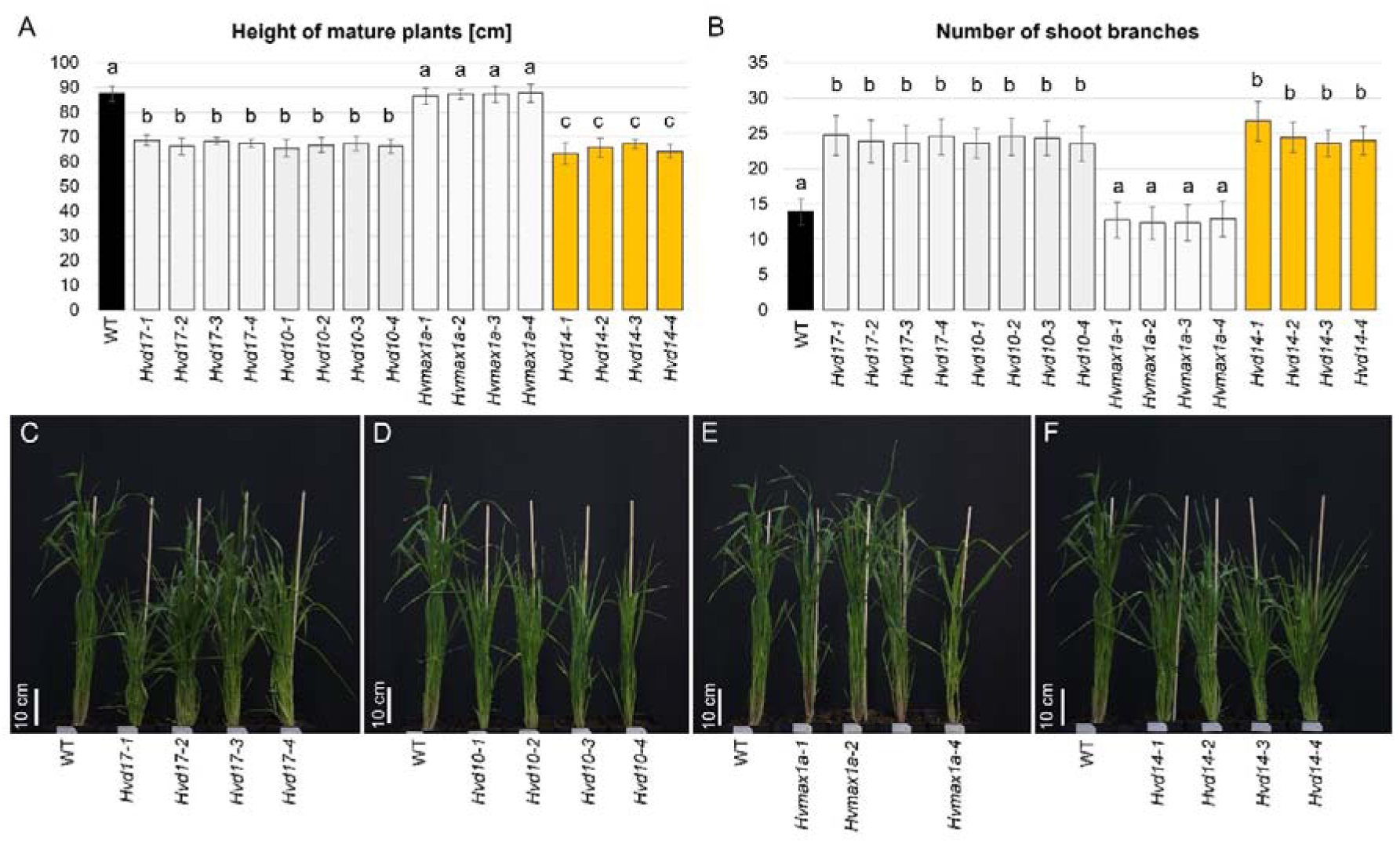
Genotype-dependent variation in plant height and shoot branching in barley SL mutants under control conditions. (A) Plant height of mature plants: WT (Golden Promise) and independent alleles of *Hvd17*, *Hvd10*, *Hvmax1a* (SL biosynthesis mutants) and *Hvd14* (SL signalling mutant). (B) Quantification of shoot branching in the same set of genotypes. (C-F). Representative two-month-old plants of WT and independent alleles of *Hvd17* (C), *Hvd10* (D), *Hvmax1a* (E), and *Hvd14* (F), illustrating genotype-specific shoot architecture. Data represent mean values ± SD (n=10 plants per genotype). Different letters indicate statistically significant differences according to one-way ANOVA followed by Tukey HSD test (p < 0.05). Scale bars = 10 cm.

Based on previous observations that the TILLING-derived barley *Hvd14.d* mutant produces smaller grains than its parent cultivar (Kelly *et al*., 2025), we conducted a detailed grain morphometric analysis across the SL mutant collection. The analysis revealed clear differences between SL mutants and the WT. Thousand-grain weight (TGW) was significantly lower in the high-tillering *Hvd17*, *Hvd10*, and *Hvd14* mutants, while the *Hvmax1a* lines did not differ from the WT (Supplementary Fig. S2). A similar pattern was seen for grain size. Both grain length and width were generally reduced in SL mutants, with the most notable decreases observed in *Hvd10* alleles, whereas *Hvmax1a* lines again resembled the WT (Supplementary Fig. S2).

### Exogenous SL application uncovers differences between biosynthesis and signalling mutants

The response of all genotypes at the seedling stage to exogenous SL application was then examined. Under control conditions, mutations in the SL biosynthesis genes *HvD17* and *HvD10*, as well as in *HvD14*, led to a significant increase in tiller number compared to WT, whereas mutants in one of the barley *MAX1* orthologues did not differ significantly from WT (Fig. 2), similar to the results in mature plants. Importantly, the tillering phenotype was consistent across all independently generated alleles within each locus, confirming a gene-specific and reproducible effect of the introduced mutations.

**Figure 2.**
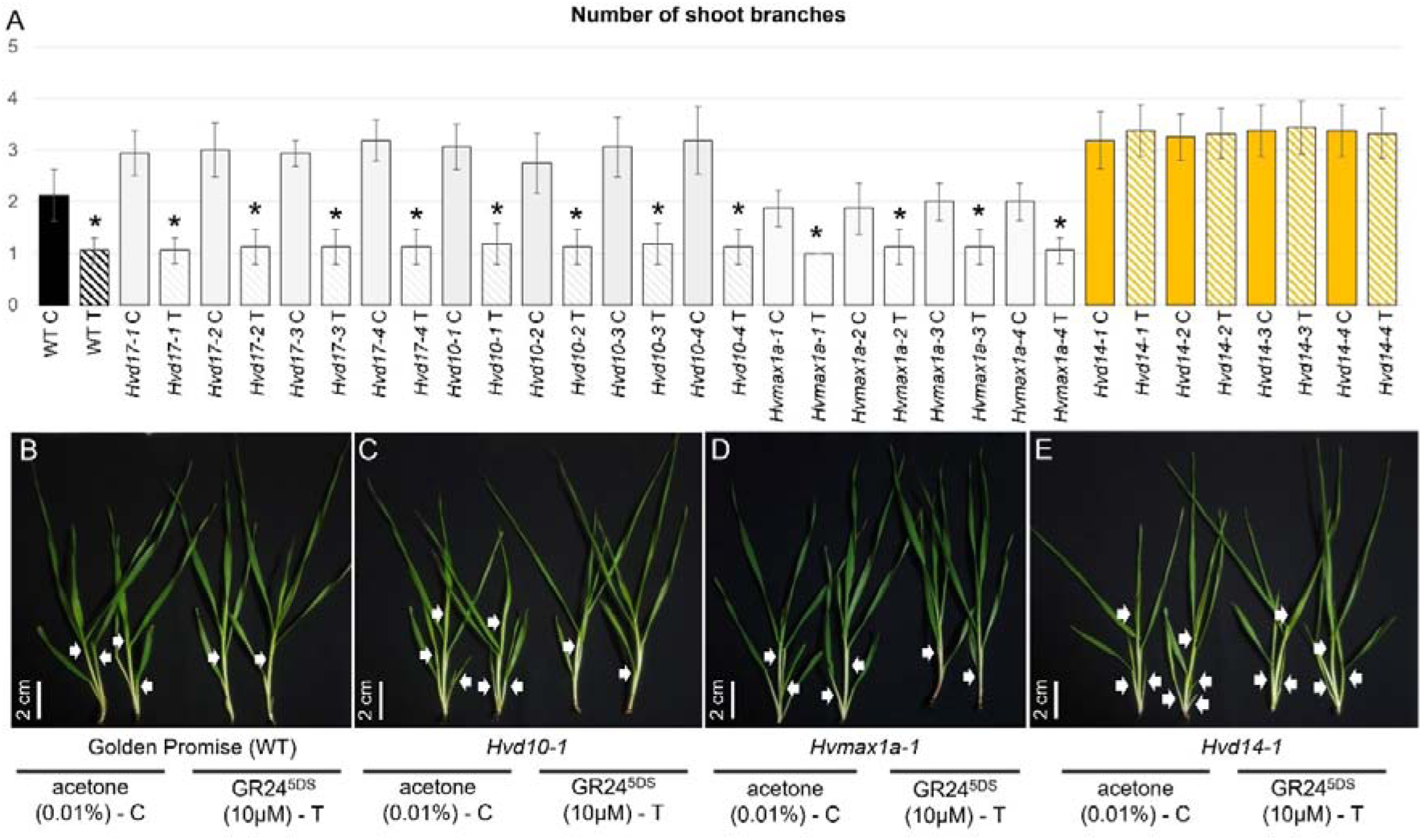
Effect of exogenous SL application on tillering in barley SL mutants. (A) Quantification of tiller number in WT barley and independent alleles of *Hvd17*, *Hvd10*, *Hvmax1a*, and *Hvd14* under control conditions (0.01% acetone; C) and following treatment with the synthetic SL analogue GR24^5DS^ (10 µM; T). (B-E) Representative seedlings of WT (B), *Hvd10-1* (C), *Hvmax1a-1* (D), and *Hvd14-1* (E) grown under control conditions or treated with GR24^5DS^. Data represent mean values ± SD (n = 16 plants per genotype and condition). Arrows indicate tillers emerging from basal nodes. Asterisks indicate statistically significant differences between control and treated plants within each genotype, as determined by a two-tailed Student’s t-test (*p* < 0.05). Scale bars = 2 cm.

Application of the synthetic SL analogue GR24^5DS^ notably decreased tiller number in WT seedlings, demonstrating that exogenous SL effectively suppresses shoot branching during early development. Consistent with previous reports based on the TILLING-derived *Hvd14.d* mutant (Daszkowska-Golec *et al*., 2023; Marzec *et al*., 2016), tillering in all analysed Cas9-triggered *Hvd14* mutants was insensitive to SL treatment, indicating impaired SL perception (Fig. 2). Conversely, SL application consistently led to a reduction of tillering across all independent alleles of the biosynthesis mutants (*Hvd17, Hvd10,* and *Hvmax1a*). The extent of tiller reduction in treated plants was comparable to that in the WT, suggesting that exogenous SL effectively compensates for endogenous SL deficiency.

### Transcriptomic alterations in SL mutants under control conditions

RNA-seq analysis was conducted on leaf tissue of four-week-old barley plants. Since independently generated alleles within each locus showed highly consistent phenotypes, a single representative line per gene was chosen for downstream analysis. The transcriptomes of these selected mutants were then compared with WT (Fig. 3, Supplementary Table S2). In the *Hvd17-1* mutant, a total of 3,313 differentially expressed genes (DEGs; log□FC ≥ 1 or ≤ -1, adjusted P ≤ 0.01) were identified relative to WT, with 2,290 genes up-regulated and 1,023 down-regulated. A similar transcriptional response was observed in *Hvd10-1*, where 3,610 DEGs were detected, including 2,074 up-regulated and 1,536 down-regulated genes. Conversely, the *Hvmax1a-1* mutant showed a significantly lower number of transcriptional changes, with 1,470 DEGs identified, among which 1,069 were up-regulated and 401 down-regulated. The *Hvd14-1* signalling mutant displayed an intermediate response, with 2,694 DEGs compared to WT, comprising 1,551 up-regulated and 1,143 down-regulated genes (Fig. 3, Supplementary Table S2).

**Figure 3.**
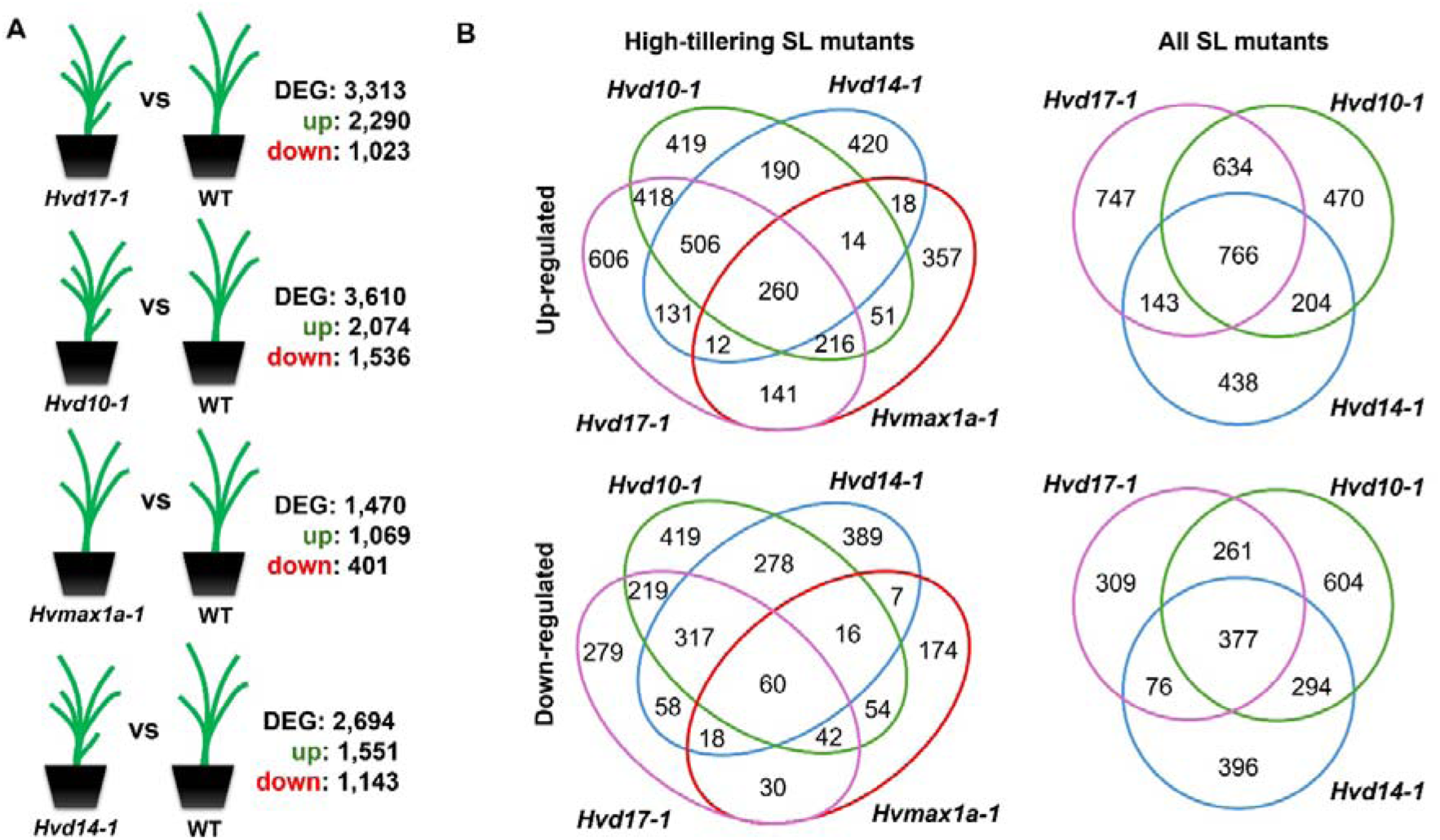
DEGs in SL mutants under control conditions. (A) Number of DEGs identified by RNA-seq in leaf tissue of four-week-old barley plants carrying mutations in *HvD17*, *HvD10*, *HvMAX1a,* and *HvD14*, each compared with the WT. For each genotype, the total number of DEGs and the numbers of up- and down-regulated genes are shown. (B) Venn diagram illustrating the overlap of up- and down-regulated DEGs among SL mutants.

A comparison of DEGs among the SL mutants revealed significant differences in the extent of transcriptomic overlap between the genotypes. When considering all four mutants together, only a small core set of genes was commonly regulated. Specifically, 260 up-regulated genes and 60 down-regulated genes were shared, corresponding to 11.4% and 5.9% of the up- and down-regulated gene sets in *Hvd17-1*, respectively, and an even smaller fraction in the other genotypes. Excluding *Hvmax1a-1* from the analysis and focusing on mutants with enhanced tillering (*Hvd17-1*, *Hvd10-1*, *Hvd14-1*) greatly increased the overlap of DEGs. Among up-regulated genes, 766 DEGs were shared by all three mutants, representing 33.5% of up-regulated genes in *Hvd17-1*, 36.9% in *Hvd10-1,* and 49.4% in *Hvd14-1*. Likewise, 377 down-regulated genes were commonly identified, accounting for 36.8% of down-regulated genes in *Hvd17-1*, 24.5% in *Hvd10-1,* and 33.0% in *Hvd14-1* (Fig. 3, Supplementary Table S3). Pairwise comparisons further indicated a particularly strong similarity between *Hvd17-1* and *Hvd10-1* mutants. These mutants shared 634 up-regulated genes, representing 27.7% of *Hvd17-1*’s up-regulated DEGs and 30.6% of *Hvd10-1’s*, as well as 261 down-regulated genes, accounting for 25.5% and 17.0% of their respective down-regulated DEGs. This constituted the largest pairwise overlap observed among all genotype combinations (Fig. 3).

Gene Ontology (GO) Biological Process enrichment analysis of shared DEGs revealed distinct patterns between high-tillering SL mutants and the full set of SL mutants. Among up-regulated DEGs shared by high-tillering mutants, the most significantly enriched terms related to amino sugar and aminoglycan catabolic processes, cell wall macromolecule catabolism, glutathione metabolism, and responses to stimuli. In contrast, the down-regulated DEGs shared by high-tillering mutants were predominantly enriched for terms associated with regulation of transcription, RNA biosynthetic processes, and gene expression. Analysis of DEGs shared by all SL mutants identified a smaller set of enriched biological processes. Up-regulated DEGs were enriched for amino acid catabolic and metabolic processes as well as responses to abscisic acid and other chemical stimuli, whereas down-regulated DEGs showed strong enrichment for regulation of transcription, RNA metabolic processes, and regulation of gene expression (Supplementary Table S4).

Transcription factor (TF) enrichment analysis was carried out using promoter sequences (1500 bp) of genes shared among SL mutants. In the DEG set common to the three high-tillering mutants (1,143 genes), 12,115 predicted TF-target regulations were identified, connecting 191 TFs to 918 target genes; among these, 82 TFs had significantly over-represented targets within the input gene set (cutoff *P* ≤ 0.05). Conversely, analysis of the DEG set shared by all four SL mutants (320 genes) revealed 3,292 regulations linking 190 TFs to 254 target genes, with 55 TFs exhibiting significant target over-representation, reflecting a more limited shared regulatory network (Supplementary Table S5). Transcription factor (TF) enrichment analysis was performed using promoter sequences (1500 bp) of genes shared between SL mutants. In the DEG set shared by the three high-tillering mutants (1,143 genes), 12,115 predicted TF-target regulations were identified, linking 191 TFs to 918 target genes; among these, 82 TFs possessed significantly over-represented targets within the input gene set (cutoff *P* ≤ 0.05). In contrast, analysis of the DEG set shared by all four SL mutants (320 genes) yielded 3,292 regulations connecting 190 TFs to 254 target genes, with 55 TFs showing significant target over-representation, consistent with a more restricted shared regulatory network (Supplementary Table S5).

Comparison of enriched TFs between the two analyses revealed 46 TFs common to both DEG sets. Within this shared TF core, several regulators acted as high-degree hubs in the high-tillering DEG set, including HORVU.MOREX.r2.2HG0148630.1 (438 target genes among the 1,143 shared DEGs), HORVU.MOREX.r2.5HG0437620.1 (395), HORVU.MOREX.r2.3HG0205590.1 (274), HORVU.MOREX.r2.4HG0287500.1 (270), and HORVU.MOREX.r2.2HG0135690.1 (255). The same TFs also ranked among the most connected regulators in the four-mutant core DEG set, with HORVU.MOREX.r2.2HG0148630.1 (111 targets) and HORVU.MOREX.r2.5HG0437620.1 (109) remaining the top hubs, indicating conservation of highly connected regulators across both shared DEG sets (Supplementary Table S5).

It is noteworthy that, although most of the key TFs did not show differential expression in the leaf tissue itself, 8 of the 46 common TFs showed differential expression in at least one mutant. This included a NAC TF (HORVU.MOREX.r2.4HG0315810.1) that was up-regulated in all four mutants (log□FC = 1.97 in *Hvd17-1*, 2.49 in *Hvd10-1*, 1.92 in *Hvmax1a-1,* and 1.78 in *Hvd14-1*), an ERF TF (HORVU.MOREX.r2.3HG0245410.1) that was induced in the three high-tillering SL mutants (log□FC = 1.40, 2.08, and 1.01 in *Hvd17-1*, *Hvd10-1,* and *Hvd14-1*, respectively), and several TFs consistently down-regulated across the high-tillering genotypes. Among the TFs that appeared in both analyses, the most highly interconnected regulators did not show any differences in expression themselves, whereas only a smaller subset of TFs showed altered expression in SL mutants (Supplementary Table S5).

### Phytohormonal profiles of SL mutants under control conditions

Phytohormone profiling showed that disrupting the SL pathway in the WT background notably impacts specific steps in the abscisic acid (ABA) and jasmonate metabolic pathways. A significant accumulation of the secondary ABA catabolite dihydrophaseic acid (DHPA) was seen in the high-tillering SL mutants Hv*d17-1*, *Hvd10-1*, and *Hvd14-1*. For phaseic acid (PA), significant increases were limited to the *Hvd17-1* and *Hvd10-1* biosynthesis mutants. Interestingly, though numerical fluctuations occurred, free ABA levels did not differ significantly across the tested lines compared to WT (Supplementary Table S6, Supplementary Fig. S3). Concerning other hormone groups, a notable reduction in the active jasmonate JA-isoleucine (JA-Ile) was observed in the high-tillering SL mutants. Conversely, the *Hvmax1a-1* mutant showed no significant changes in any of the measured hormone levels, consistent with its wild-type-like branching phenotype (Supplementary Fig. S3). Importantly, despite evident morphological differences in tillering, no statistically significant changes were detected in the steady-state levels under these conditions of active cytokinins (tZ, cZR), auxins (IAA), or gibberellins (GAs) in the shoots of any mutant line (Supplementary Table S6).

### Drought-induced physiological responses of SL mutants

Based on the previous evidence of increased drought sensitivity in the barley TILLING mutant *Hvd14.d* (Daszkowska-Golec *et al*., 2023; Marzec, Daszkowska-Golec, *et al*., 2020), along with reports from other plant species showing that disruption of SL biosynthesis or signalling undermines drought tolerance (Haider *et al*., 2018; Li *et al*., 2020), we analysed drought-induced physiological responses in a collection of SL mutants. Drought stress caused a significant reduction in biomass across all genotypes (Fig. 4). WT plants retained about 35% of the dry weight of the control group, while all SL mutants were more severely affected. The three high-tillering SL mutants (*Hvd17-1*, *Hvd10-1*, and *Hvd14-1*) exhibited a strongly uniform response, with dry mass reduced to approximately 22-23% of control values. Conversely, the *Hvmax1a-1* mutant showed a less marked reduction, retaining roughly 26% of control dry mass, indicating a partial decrease in drought sensitivity compared to the high-tillering SL genotypes (Fig. 4).

**Figure 4.**
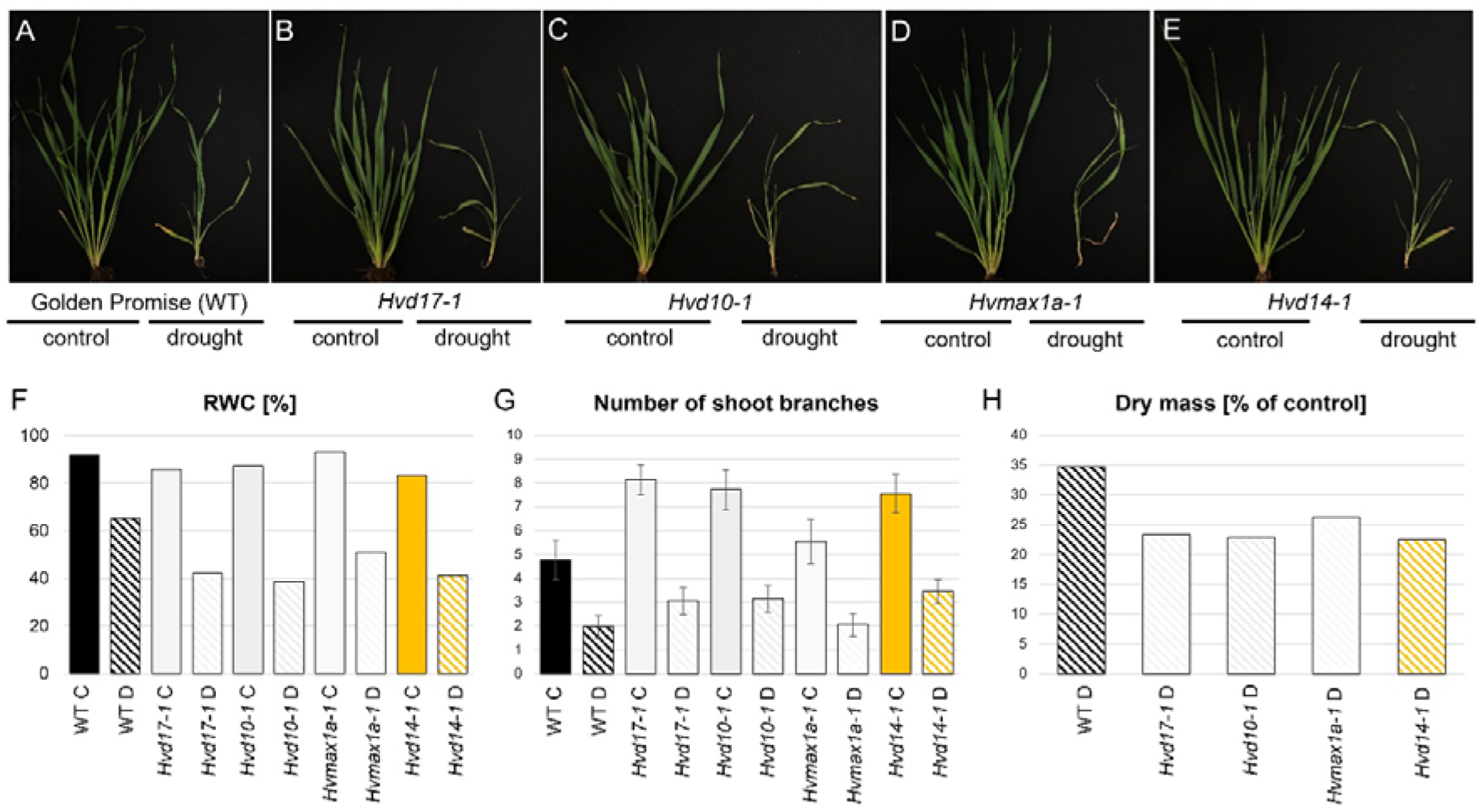
Physiological responses to drought in barley SL mutants. (A-E) Representative phenotypes of WT and SL mutants (*Hvd17-1, Hvd10-1, Hvmax1a-1,* and *Hvd14-1*) grown under control and drought conditions. (F) Relative water content (RWC), (G) number of shoot branches per plant, and (H) dry mass expressed as a percentage of control conditions for all analysed genotypes.

Changes in biomass accumulation were accompanied by fluctuations in the plants’ water status. Under drought conditions, relative water content (RWC) decreased in all genotypes, but the decline was consistently more pronounced in SL mutants than in WT plants. WT leaves maintained an RWC of 65.0%, whereas the three high-tillering SL mutants showed clearly lower, very similar values: 42.2% in *Hvd17-1*, 38.6% in *Hvd10-1*, and 41.4% in *Hvd14-1*. The *Hvmax1a-1* mutant displayed an intermediate response, with a higher RWC (50.9%) than the high-tillering SL mutants, but still notably lower than WT (Fig. 4).

Drought stress also had a significant effect on shoot branching, with genotype-specific responses that, under control conditions, depended on the initial tillering level. In the control treatment, notable differences in shoot branching were observed among genotypes, with the highest branching in the high-tillering SL mutants (8.14, 7.73, and 7.55 branches per plant in *Hvd17-1*, *Hvd10-1*, and *Hvd14-1*, respectively) and lower values in the WT (4.77) and *Hvmax1a-1* (5.55). Under drought stress, the number of shoot branches decreased across all genotypes; however, the relative extent of this reduction was similar within each genetic background. WT plants reduced the number of shoot branches from 4.77 to 1.95, a 41% reduction relative to the control. A comparable proportional reduction was seen in *Hvd17-1* (3.05 vs 8.14; 37% of control), *Hvd10-1* (3.14 vs 7.73; 41%), and *Hvmax1a-1* (2.05 vs 5.55; 37%). The *Hvd14-1* mutant exhibited a slightly weaker reduction, with 3.45 branches remaining compared to 7.55 in control conditions, representing 46% of the control level (Fig. 4).

To further characterise the physiological basis of the differential drought responses observed among SL mutants, we analysed photosystem II (PSII) performance. In WT plants, drought stress caused a moderate reduction in φP□ (Fv/Fm) and a significant decline in the performance index PI ABS, indicating impaired overall PSII functionality. These changes were accompanied by increased ABS/RC, reflecting a reduction in the number of active PSII reaction centres, and elevated DI□/RC, indicating enhanced dissipation of excess excitation energy, a characteristic of drought-induced photoprotective adjustments in PSII (Supplementary Fig. S4). The three high-tillering SL mutants displayed largely similar but quantitatively different PSII responses to drought. In *Hvd10-1* and *Hvd17-1*, drought caused reductions in φP□ and significant decreases in PI ABS exceeding those in WT plants. Both mutants also showed marked increases in ABS/RC and DI□/RC, indicating pronounced reaction centre deactivation and increased energy dissipation. These patterns suggest that a limited capacity to maintain PSII efficiency under water deficit might be related to their substantial reductions in biomass accumulation and leaf water status. Among the high-tillering SL mutants, *Hvd14-1* showed the most severe PSII impairment under drought conditions. This genotype exhibited the strongest decline in both φP□ and PI ABS, along with the highest increases in ABS/RC and DI□/RC. These changes suggest a significant loss of functional PSII reaction centres and the possibility that non-photochemical quenching (NPQ) processes are dominant, a finding partially reflected by an increased DI□/RC ratio, consistent with pronounced photoinhibitory stress. Unlike the high-tillering SL mutants, *Hvmax1a-1* displayed a notably more stable PSII response to drought. Although drought stress reduced PI ABS in this genotype, the decline was smaller than in WT and considerably lower than in the high-tillering SL mutants. Similarly, increases in ABS/RC and DI□/RC were more limited, indicating better preservation of functional reaction centres and more efficient regulation of excitation energy under water deficit. The comparatively minor reduction in φP□ further suggests that PSII photochemistry in *Hvmax1a-1* is less susceptible to drought-induced inhibition (Supplementary Fig. S4).

### Drought-induced transcriptome responses of SL mutants

To characterise transcriptional responses to drought, RNA-seq analysis was conducted to compare drought-treated and control plants within each genotype. In WT plants, drought stress led to the identification of 8,700 DEGs (log□FC ≥ 1 or ≤ -1, adjusted P ≤ 0.01), including 4,848 up-regulated and 3,852 down-regulated genes (Fig. 5, Supplementary Table S7). A considerably stronger response was observed in the high-tillering SL mutants. In *Hvd17-1*, a total of 13,056 DEGs were detected, comprising 6,468 up-regulated and 6,588 down-regulated genes, representing the most extensive transcriptional reprogramming across all analysed genotypes. Similarly, *Hvd10-1* exhibited 12,347 DEGs, with 5,990 genes up-regulated and 6,357 down-regulated under drought conditions. The signalling mutant *Hvd14-1* showed a similarly strong response, with 10,964 DEGs, including 5,709 up-regulated and 5,255 down-regulated genes. In contrast, the SL biosynthesis mutant *Hvmax1a-1* displayed a significantly reduced transcriptional response, with 8,233 DEGs identified (3,357 up-regulated and 4,876 down-regulated), suggesting decreased transcriptional sensitivity to drought (Fig. 5, Supplementary Table S7).

**Figure 5.**
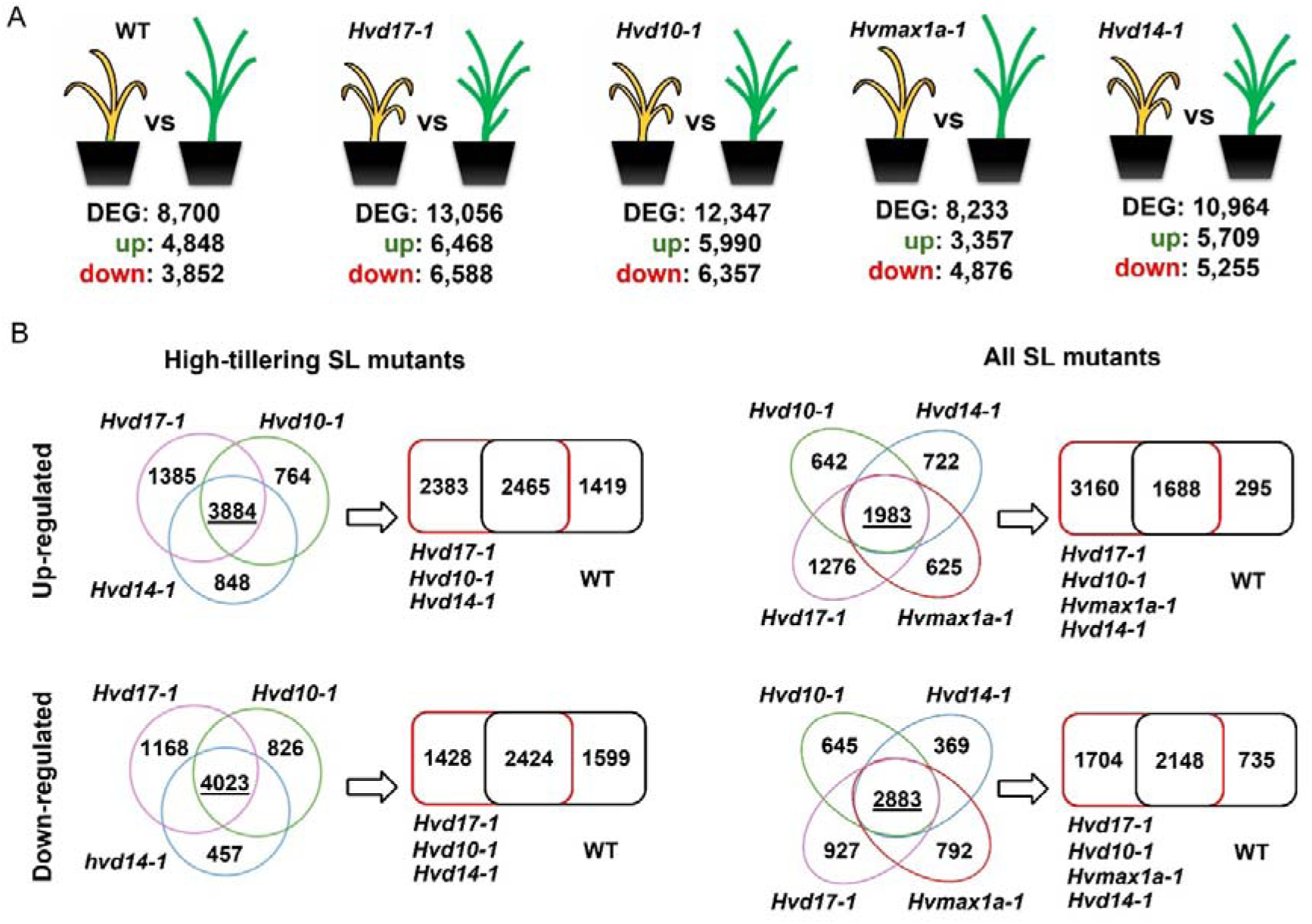
Drought-induced transcriptomic responses in SL mutants. (A) Number of DEGs identified by RNA-seq in barley plants with mutations in *HvD17*, *HvD10*, *HvMAX1a,* and *HvD14* following drought treatment, each compared with the respective control condition within the same genotype. For each genotype, the total number of DEGs and the counts of up-and down-regulated genes are displayed. (B) Venn diagrams showing the overlap of drought-responsive up- and down-regulated DEGs among the three high-tillering SL mutants (*Hvd17-1*, *Hvd10-1,* and *Hvd14-1*) and across all four SL mutants. For each comparison, the number of shared genes among mutants and their overlap with the WT drought response are indicated.

The drought-responsive DEGs were grouped into genes common to all four SL mutants or to the three high-tillering SL genotypes, and into genes specific to either mutant groups or WT plants under drought conditions (Fig. 5, Supplementary Table S8). When considering all four SL mutants, the shared drought-responsive core included 1,983 up-regulated and 2,883 down-regulated genes. This four-mutant core made up 30.7-34.7% of the up-regulated genes in the high-tillering SL mutants (*Hvd17-1* 30.7%, *Hvd10-1* 33.1%, *Hvd14-1* 34.7%), but a significantly larger portion of the *Hvmax1a-1* up-regulated set (59.1%). A similar trend was seen for down-regulated genes, where the four-mutant core accounted for 43.8-54.9% of down-regulated genes in the high-tillering SL mutants (*Hvd17-1* 43.8%, *Hvd10-1* 45.4%, *Hvd14-1* 54.9%), and 59.1% in *Hvmax1a-1* (Fig. 5, Supplementary Table S8).

Consistent with this, most genes in the four-mutant core overlapped with the WT drought response. Specifically, 1,688 up-regulated genes (85.1%) and 2,148 down-regulated genes (74.5%) were also drought-responsive in WT. Only 295 up-regulated and 735 down-regulated genes were unique to the four-mutant core, suggesting that the intersection across all four mutants primarily reflects the canonical WT drought programme. In contrast, focusing on the three high-tillering SL mutants significantly expanded the shared drought-responsive set. The high-tillering core included 3,884 up-regulated and 4,023 down-regulated genes, representing 60.0-68.0% of up-regulated genes (*Hvd17-1* 60.0%, *Hvd10-1* 64.8%, *Hvd14-1* 68.0%) and 61.1-76.6% of down-regulated genes (*Hvd17-1* 61.1%, *Hvd10-1* 63.3%, *Hvd14-1* 76.6%). Importantly, this expanded core showed only partial overlap with the WT drought response: 2,465 up-regulated genes (63.5%) and 2,424 down-regulated genes (60.3%) were shared with WT, while a significant proportion was specific for high-tillering mutants (1,419 up-regulated and 1,599 down-regulated genes not detected in WT) (Fig. 5, Supplementary Table S9).

GO Biological Process enrichment analysis of genotype-specific drought-responsive DEGs revealed clear functional differences between WT and SL mutants. Among up-regulated genes specific to the three high-tillering SL mutants, enriched terms predominantly related to the cell cycle and genome maintenance, including chromosome condensation, mitotic cell cycle processes, DNA replication, and DNA repair. In contrast, down-regulated genes specific to high-tillering SL mutants mainly associated with photosynthesis, phenylpropanoid metabolism, carbohydrate metabolism, and defence-related processes (Supplementary Table S10). Analysis of WT-specific drought-responsive genes relative to high-tillering SL mutants showed a distinct enrichment profile. WT-specific up-regulated genes were enriched for phenylpropanoid and lignin metabolism, translation-related processes, and biosynthetic pathways, whilst WT-specific down-regulated genes were dominated by terms related to regulation of transcription, RNA biosynthesis, and gene expression (Supplementary Table S10). When genotype-specific DEGs were defined in comparison to all four SL mutants, the clustering of biological processes was less pronounced. Up-regulated genes specific to all SL mutants were primarily associated with DNA metabolism and cell cycle processes, whilst down-regulated genes were enriched in photosynthesis and phosphorylation pathways. Conversely, WT-specific genes relative to all SL mutants showed enrichment for oxylipin and phenylpropanoid metabolism among up-regulated genes, and regulation of transcription and defence responses among down-regulated genes (Supplementary Table S10).

In the WT-specific drought-responsive gene set, 41,937 predicted TF-target regulations were identified, linking 191 TFs to 2,927 target genes, among which 96 TFs showed significantly over-represented targets within the input gene set (cutoff *P* ≤ 0.05). In contrast, the mutant-specific set of drought-responsive genes yielded 31,675 predicted regulations connecting the same 191 TFs to 2,236 target genes, with 77 TFs displaying significant target over-representation, indicating a reduced regulatory complexity in SL mutants under drought conditions. Analysis of drought-responsive genes shared between WT and SL mutants revealed 87,347 predicted TF-target regulations involving 191 TFs and 6,257 target genes, among which 97 TFs had significant target over-representation (Supplementary Table S11), defining a substantial SL-independent core regulatory network. A comparative analysis of TF enrichment profiles across WT-specific, mutant-specific, and shared drought-responsive gene sets further identified 20 TFs that were uniquely enriched in the WT-specific response. Notably, 13 of these TFs were also WT-specific when compared against all four SL mutants, and five TFs were themselves differentially expressed in WT (Table 2).

### Phytohormonal responses of SL mutants under drought conditions

Drought treatment triggered a pronounced activation of the ABA pathway across all analysed genotypes. In WT plants, a significant accumulation of ABA was accompanied by strong increases in its major catabolites and conjugates, including PA, DHPA, ABA-glucose ester (ABAGlc), and ABA-glucoside (ABAGlu). A similar pattern was observed in all SL mutants, indicating a conserved drought-induced activation of ABA metabolism. Notably, the high-tillering SL mutants *Hvd17-1, Hvd10-1,* and *Hvd14-1* showed significant increases in multiple ABA metabolites, particularly DHPA and ABAGlc, whereas the rise in free ABA was more variable across genotypes. Despite its WT-like branching phenotype, *Hvmax1a-1* also showed significant accumulation of ABA derivatives under drought, confirming that perturbation of SL biosynthesis affects ABA turnover even in the absence of a strong architectural phenotype (Supplementary Table S12, Fig. 6).

**Figure 6.**
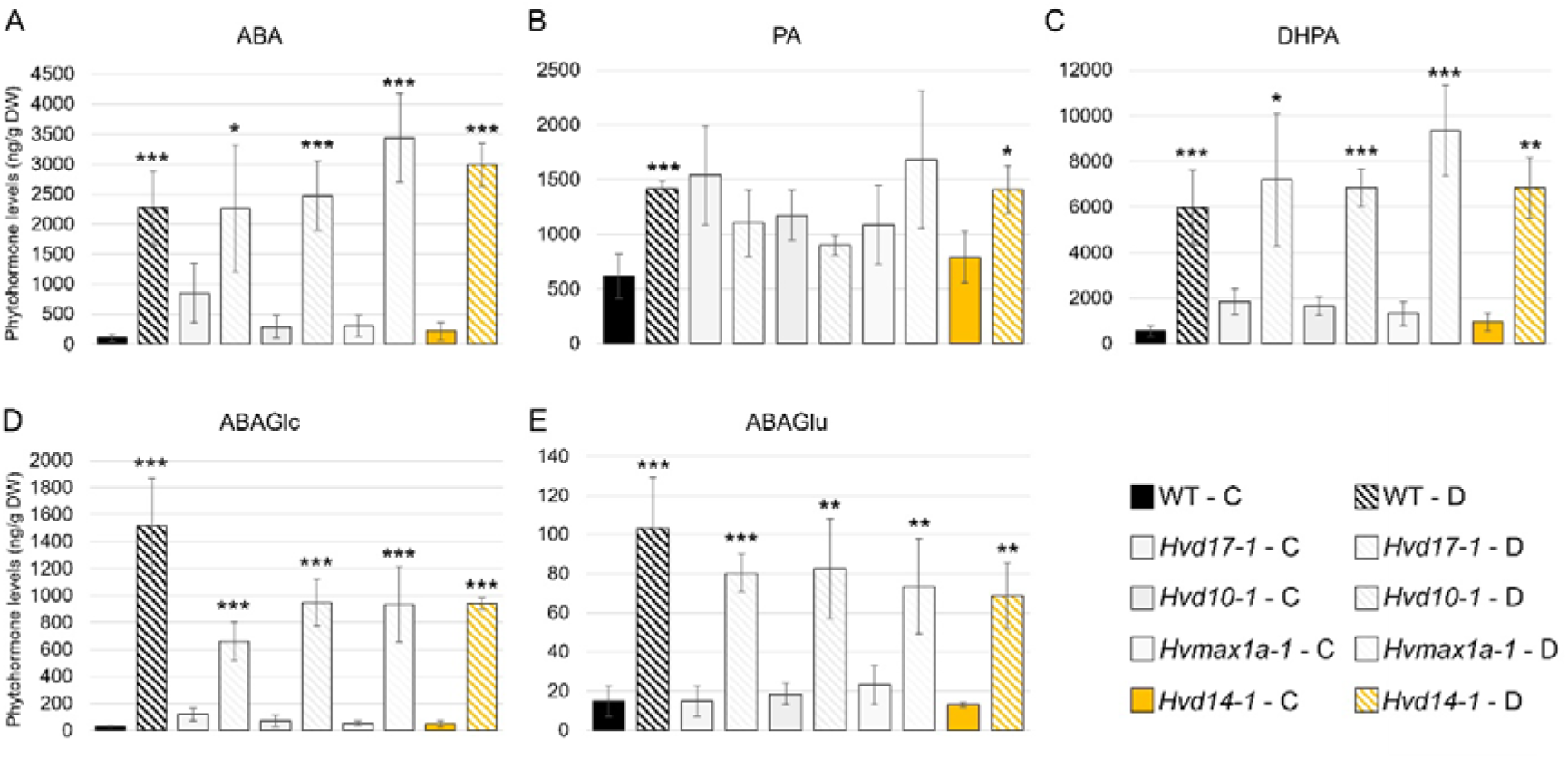
Drought-induced activation of ABA metabolism in barley SL mutants. (A) Abscisic acid (ABA), (B) phaseic acid (PA), (C) dihydrophaseic acid (DHPA), (D) ABA-glucose ester (ABAGlc), and (E) ABA-glucoside (ABAGlu) levels in WT and SL mutants under control (C) and drought (D) conditions. Data represent mean values ± SE (n = 4). Asterisks indicate statistically significant differences between control and drought within a given genotype according to Student’s *t*-test (*p* < 0.05, *p* < 0.01, *p* < 0.001). Phytohormone concentrations are expressed as ng g□¹ dry weight (DW).

Beyond the ABA pathway, drought triggered additional, more genotype-specific hormonal adjustments. Significant decreases in selected gibberellin intermediates were observed in several mutant lines, while changes in jasmonates were limited and depended on genotype. Notably, the strongly tillering mutants did not show any consistent drought-induced shifts in steady-state auxin or active cytokinin pools (Supplementary Table S12), suggesting that their contrasting branching phenotype under stress cannot be attributed to major global changes in these hormone classes.

## Discussion

### Architectural phenotypes uncover functional diversity within the barley SL pathway

The architectural phenotypes observed in this study reinforce the conserved role of SLs as central regulators of shoot branching and plant stature (Dun *et al*., 2023). Mutations affecting early SL biosynthesis (HvD17/CCD7 and HvD10/CCD8) and SL perception (HvD14) produced a highly consistent phenotype of enhanced tillering and reduced height, in line with the canonical SL-deficient syndrome described across monocots and dicots (Gomez-Roldan *et al*., 2008; Umehara *et al*., 2008). In barley, this architectural response has previously been demonstrated in the TILLING-derived *Hvd14.d* mutant, confirming that disruption of SL perception alone is sufficient to trigger strong branching and semi-dwarfism (Marzec *et al*., 2016). Similar phenotypes have also been described for mutations affecting later stages of biosynthesis, including those in the *HvLBO* gene, which increase tillering despite retaining detectable SL levels in roots and exudates. This suggests a functional specialization among SL molecules with distinct physiological roles (Inoue *et al*., 2025).

Unlike the marked phenotype of *Hvd17, Hvd10*, and *Hvd14* mutants, disrupting *HvMAX1a* did not affect plant height or tillering, indicating functional divergence within the downstream SL biosynthetic pathway. These results imply that enzymes functioning downstream of carlactone mainly influence SL structural diversity rather than overall hormone levels (Ou *et al*., 2026). In monocots, multiple CYP711A/MAX1 paralogues catalyse different oxidative reactions, producing a variety of canonical and non-canonical SLs with potentially specialised biological functions. Evidence for this diversification comes from studies showing that various CYP711A/MAX1 paralogues catalyse distinct oxidative steps, creating different SL structures with potentially unique roles (Cardoso *et al*., 2014; Yoneyama *et al*., 2018; Zhang *et al*., 2014). For instance, rice has five MAX1 homologues that act at different stages of SL biosynthesis, including converting carlactonoic acid into 4-deoxyorobanchol and orobanchol, demonstrating how pathway diversification leads to structurally varied SLs (Marzec, Situmorang, *et al*., 2020). A similar duplication and diversification of *MAX1* genes have been observed in *Triticum aestivum* (wheat), where 13 MAX1 homologues form several clades with distinct tissue-specific expression, with some predominating in roots and others in aerial parts, indicating a partitioning of SL biosynthesis (Sigalas *et al*., 2023). This functional diversification explains the *Hvmax1a* phenotype observed here and highlights the importance of considering SL structural diversity when analysing mutant effects on plant architecture.

### Effects of SL disruption on grain development

Disruption of SL signalling clearly impacts grain development and establishes a link between architectural traits and reproductive performance. Consistent with detailed analyses of the TILLING-derived *Hvd14.d* receptor mutant, SL-insensitive barley plants produced more tillers and grains, but individual grains were smaller. This trade-off largely offset potential yield increases associated with greater spike production (Kelly *et al*., 2025). This phenotype remained stable across environments and persisted even when tiller number was experimentally reduced, indicating that the reduction in grain size cannot be explained solely by altered assimilate availability but also reflects a direct role of SL signalling in regulating grain filling and sink strength (Kelly *et al*., 2025). Importantly, evidence from other cereals supports a conserved role of SLs in reproductive development. It’s been shown that in *Zea mays* (maize), SL-regulated transcriptional networks directly contribute to the determination of grain size and its evolutionary diversification (Guan *et al*., 2023). In line with classical crop physiology concepts suggesting that grain weight is predominantly sink-limited (Alvarez Prado *et al*., 2013), these findings indicate that SL signalling coordinates whole-plant resource allocation with local developmental regulation of grain growth. Within this framework, the lack of a strong phenotype in *Hvmax1* mutant grains in our study likely reflects functional redundancy among *MAX1* paralogues or the differential contribution of individual SL intermediates to reproductive signalling. Overall, our observations position SLs as key coordinators of the architecture-yield relationship, functioning at both the whole-plant and local levels through resource distribution and regulation of grain development.

### Genotype- and context-dependent hormonal signatures of SL signalling disruption

Hormonal differences between SL mutants closely reflected their architectural phenotypes. The three high-tillering mutants (*Hvd17, Hvd10,* and *Hvd14*) showed consistent changes in ABA turnover and jasmonate signalling under control conditions, whereas the *Hvmax1a* mutant displayed a largely WT-like hormonal profile. This distinction implies that disruption of core SL biosynthesis or perception causes coordinated hormonal reprogramming, while perturbing a single *MAX1* paralogue is insufficient to significantly alter steady-state hormone levels. Comparing phytohormone profiles between the genome editing-derived SL mutants analysed here and the previously characterised TILLING mutant *Hvd14.d* reveals a strong context-dependent hormonal response to SL disruption. In the *Hvd14.d* mutant (cv. Sebastian background), four-week-old plants showed notable differences: wild-type accumulated higher levels of ABA, JA, and SA, while the mutant had increased cytokinin levels (Korek, Mehta, *et al*., 2025). Conversely, our analyses in the cv. Golden Promise background detected more moderate changes under control conditions, primarily impacting ABA turnover rather than free ABA levels. These discrepancies likely stem from both methodological and biological factors. Chemical mutagenesis had caused thousands of unknown background mutations in the *Hvd14.d* line of cv. Sebastian, some of which may be responsible for specific features and responses. In the comparison of the mutants, it should also be taken into account that hormone measurements in *Hvd14.d* were performed on fresh whole-shoot tissue, whereas our new dataset is based on dry leaf material, which can influence the detection of mobile and conjugated hormone pools (Almeida Trapp *et al*., 2014). Additionally, cultivar-specific hormone homeostasis may contribute to the differing profiles. The cv. Sebastian background contains the widely used semi-dwarfing allele *sdw1/denso*, which involves mutations in the *HvGA20ox2* gene encoding GA 20-oxidase, a key enzyme in gibberellin biosynthesis (Jia *et al*., 2009). Reduced GA production linked to *sdw1* alters plant stature and affects hormonal crosstalk involving cytokinins and ABA (Cheng *et al*., 2023), potentially reshaping the downstream effects of SL signalling disruption (Korek, Mehta, *et al*., 2025). By contrast, cv. Golden Promise carries a distinct semi-dwarfing mutation (*breviaristatum-e, ari-e*) affecting G-protein signalling rather than GA biosynthesis (Liu *et al*., 2014), indicating that the hormonal baseline underlying SL mutant phenotypes varies significantly between the two genetic backgrounds. Overall, this comparison highlights that hormonal responses to SL disruption are influenced by developmental stage, tissue context, and genetic background, supporting a model in which SL signalling interacts with flexible hormonal networks rather than exerting fixed control over individual hormone pools.

### Disruption of SL biosynthesis and signalling compromises plant water status and PSII performance under drought

This study expands on previous evidence linking SL signalling with drought tolerance in barley by showing that genome editing-driven mutants in the Golden Promise background consistently display a drought-sensitive physiological phenotype. Earlier research using the TILLING-derived *Hvd14.d* mutant demonstrated that disrupting SL perception hampers drought adaptation by impairing stomatal regulation, modifying leaf structure, and decreasing photosynthetic efficiency (Daszkowska-Golec *et al*., 2023; Marzec, Daszkowska-Golec, *et al*., 2020). Alongside findings from other species indicating heightened drought sensitivity in SL-deficient or SL-insensitive genotypes (Ha *et al*., 2014; Haider *et al*., 2018; Liu *et al*., 2015; Visentin *et al*., 2016), our results reinforce the idea of a conserved role for SL in drought acclimation.

All high-tillering SL mutants, including those affecting biosynthesis (*Hvd17, Hvd10*) and signalling (*Hvd14*), exhibited more pronounced drought-induced reductions in biomass and leaf water content than WT plants, suggesting a reduced ability to maintain water balance under water deficit. Similar physiological effects have been associated with increased water loss and impaired stomatal behaviour in SL mutants across different species (Ha *et al*., 2014; Lv *et al*., 2018; Visentin *et al*., 2016). In contrast, the *Hvmax1a* mutant displayed a different response pattern. Under normal conditions, *Hvmax1a* closely resembled WT plants, aligning with the role of MAX1 enzymes in structural diversification rather than core SL biosynthesis. However, during drought, *Hvmax1a* showed an intermediate phenotype, responding more strongly than WT but less severely than the high-tillering mutants. This suggests that while MAX1a-derived SLs are mostly unnecessary for growth under favourable conditions, they aid in stress adaptation, potentially by producing specific SL molecules involved in drought-responsive signalling. However, this remains to be tested directly. Our observations align with previous evidence that individual MAX1 homologues can differentially regulate SL composition and stress responses (Marzec, Situmorang, *et al*., 2020).

Photosynthetic analyses revealed that impaired growth and hydration in SL mutants were accompanied by reduced PSII performance. While WT plants displayed typical drought-induced photoprotective adjustments, characterised by moderate declines in φP□ and increased energy dissipation (Daszkowska-Golec *et al*., 2019), the high-tillering mutants showed stronger decreases in φP□ and PI ABS, along with pronounced increases in ABS/RC and DI□/RC. These responses indicate extensive deactivation of the reaction centre and enhanced non-photochemical energy dissipation, consistent with limited photoprotective capacity and increased photoinhibitory pressure (Filek *et al*., 2015; Oukarroum *et al*., 2016). The strongest PSII impairment observed in *Hvd14* highlights the central role of D14-mediated perception and aligns with earlier multi-omics analyses demonstrating widespread SL-dependent reprogramming of drought responses, including altered ROS homeostasis and weakened antioxidant capacity (Daszkowska-Golec *et al*., 2023; Visentin *et al*., 2016).

Our findings support a model in which SLs enhance drought tolerance by maintaining plant water status and preserving PSII functionality. Loss of SL function results in impaired hydration, increased reliance on energy dissipation, and reduced photosynthetic efficiency, ultimately limiting biomass accumulation. The similarity of phenotypes across independent mutants and genetic backgrounds further emphasises the robustness of SL-dependent drought adaptation and highlights SL signalling as a key integrator linking plant architecture with physiological resilience to water deficit.

### ABA metabolism is activated, but the ABA response seems reduced in SL mutants during drought

Drought treatment triggered a clear activation of the ABA pathway across all analysed genotypes, evidenced by the accumulation of ABA along with its major catabolites and conjugates. WT plants showed a coordinated rise in free ABA, PA, DHPA, and glucosylated forms, aligning with the well-known role of ABA turnover and reversible conjugation in modulating drought responses (Kuromori *et al*., 2022; Nambara and Marion-Poll, 2005). A broadly similar metabolic profile was observed in all SL mutants, suggesting that disruption of SL biosynthesis or signalling does not inhibit drought-induced ABA metabolism activation. However, the high-tillering mutants *Hvd17*, *Hvd10,* and *Hvd14* displayed particularly strong accumulation of DHPA and ABA-glucosylated forms, although variations in free ABA levels were more variable. This pattern indicates enhanced flux through ABA catabolic and conjugation pathways rather than a straightforward increase in steady-state ABA levels. These dynamics support the idea that ABA-glucosyl esters serve as inactive storage pools that can be rapidly mobilised during dehydration, while hydroxylation-derived metabolites reflect active hormone turnover (Nambara and Marion-Poll, 2005). Notably, the *Hvmax1a* mutant also exhibited significant accumulation of ABA derivatives despite its WT-like structure under control conditions, implying that alterations in SL composition mainly affect ABA homeostasis under stress rather than during normal growth.

Importantly, the hormonal profile observed here mirrors earlier findings for the TILLING-derived *Hvd14.d* mutant, in which drought or rapid dehydration caused similar increases in ABA and its metabolites in both WT and mutant plants (Daszkowska-Golec *et al*., 2023; Marzec, Daszkowska-Golec, *et al*., 2020). Along with the present data, these observations suggest that SL deficiency impacts not the ability to accumulate ABA, but rather the efficiency with which ABA signals are converted into adaptive physiological responses. A possible mechanistic explanation involves reduced stomatal responsiveness to ABA in SL mutants. Decreased ABA sensitivity and altered stomatal behaviour have been reported for SL-deficient or SL-insensitive genotypes across species (Ha *et al*., 2014; Liu *et al*., 2015; Marzec, Daszkowska-Golec, *et al*., 2020; Visentin *et al*., 2016), while SL itself can promote stomatal closure through both ABA-dependent and ABA-independent pathways involving reactive oxygen species and SLAC1 activation (Korek and Marzec, 2023). In this context, the strong activation of ABA metabolism seen in the high-tillering mutants may reflect a compensatory response to inadequate stomatal regulation, ultimately causing greater water loss, impaired hydration, and increased photoinhibitory stress. This model is consistent with published evidence but was not directly tested here, as stomatal measurements were outside the scope of the present study.

### SLs shape drought-responsive transcriptional networks in barley

Drought caused a larger DEG set in the high-tillering SL mutants than in WT, indicating transcriptional hyperactivation under SL deficiency. Similar transcriptome amplification has been reported in SL-deficient or SL-insensitive genotypes of other species, where increased drought sensitivity was associated with altered ABA-dependent gene expression (Ha *et al*., 2014; Liu *et al*., 2015; Visentin *et al*., 2016). Along with earlier findings for *Hvd14.d* (Daszkowska-Golec *et al*., 2023; Marzec, Daszkowska-Golec, *et al*., 2020), this supports the view that the expanded DEG sets in SL mutants mainly reflect increased stress sensitivity rather than improved acclimation. The four-mutant core largely overlaps with WT, indicating that canonical drought signalling remains SL-independent and probably reflects conserved ABA-driven responses (Shinozaki and Yamaguchi-Shinozaki, 2007). In contrast, the expanded high-tillering core, which is only partially shared with WT, reveals a mutant-specific programme characterised by repression of photosynthesis and primary metabolism alongside induction of DNA replication and repair processes, consistent with stress-driven cellular maintenance signatures described under severe drought (Fujita *et al*., 2011; Nakabayashi *et al*., 2014) and previously observed in *Hvd14.d* (Daszkowska-Golec *et al*., 2023).

Differences were also clear at the regulatory level. The lower number of enriched TFs and TF-target interactions in mutant-specific gene sets suggests simplified transcriptional control under SL deficiency, aligning with earlier reports of weakened ABA-dependent transcriptional activation in SL mutants (Daszkowska-Golec *et al*., 2023; Li *et al*., 2020). Likewise, Table 1 highlights a subset of TFs uniquely enriched in the WT drought response, several of which belong to ERF/DREB, WRKY, NAC, and MYB families previously linked to SL-dependent regulation of drought responses (Joshi *et al*., 2016; Liu *et al*., 2024; Zhang *et al*., 2022). These TF groups are well known as integrators of ABA and ROS signalling (Chen *et al*., 2025; Postiglione and Muday, 2020), and some representatives identified in Table 1 (e.g., WRKY40-, NAC- and MYB-type regulators) are recognised for controlling wax deposition, oxidative stress responses, and cell wall remodelling (Nakabayashi *et al*., 2014). Reduced activation of these pathways has been observed in *Hvd14.d* (Daszkowska-Golec *et al*., 2023; Marzec, Daszkowska-Golec, *et al*., 2020), supporting the concept that SL signalling facilitates coordinated activation of protective metabolic modules.

**Table 1.** Transcription factors uniquely enriched in the WT-specific drought response.

| TF ID | No. of targets * | TF family | Arabidopsis homolog | HORVU v2 ID (HORVU.MOREX.r2.) | log2FC (WT D vs C) | Description |
| --- | --- | --- | --- | --- | --- | --- |
| <u>**MLOC_67851**</u> | 196 | WRKY | AT2G38470 (WRKY33) | 3HG0254080 |  | Negative regulator of GA and ABA signalling involved in responses to biotic and abiotic stresses |
| <u>**MLOC_12079**</u> | 318 | WRKY | AT5G26170 (WRKY50) | 1HG0023190 |  | Regulator of JA-mediated defence signalling and stress responses |
| <u>**MLOC_4830**</u> | 42 | SBP | AT2G47070 (SPL1) | 5HG0440620 |  | TF involved in transcriptional regulation |
| **MLOC_15725** | 817 | B3 | AT3G26790 (FUS3) | 3HG0284730 |  | Regulator of flowering time and developmental transitions |
| <u>**MLOC_60958**</u> | 628 | C2H2 | AT2G02080 (IDD4) | 6HG0499980 |  | Zinc-finger transcription factor involved in transcriptional regulation |
| <u>**MLOC_36657**</u> | 171 | WRKY | AT5G13080 (WRKY75) | 2HG0096750 |  | Stress-responsive TF involved in regulation of defence-related gene |
| <u>**MLOC_14363**</u> | 551 | C2H2 | AT5G66730 (IDD12) | 4HG0332420 | -1,14 | Positive regulator of phytochrome signalling pathways |
| <u>**MLOC_56103**</u> | 125 | NAC | AT1G12260 (NAC007) | 6HG0496420 |  | Regulator of secondary cell wall biosynthesis and PCD |
| <u>**MLOC_25460**</u> | 158 | Trihelix | AT1G76890 (GT2) | .6HG0449910 | 1,14 | TF involved in regulation of gene expression during development |
| <u>**MLOC_78652**</u> | 500 | C2H2 | AT1G03840 (IDD3) | 3HG0208500 |  | Regulator of tissue boundary formation and cell division |
| **<u>MLOC_57518</u>** | 480 | C2H2 | AT1G55110<br>(IDD7) | 3HG0221400 |  | TF involved in regulation of<br>gene expression |
| **MLOC_75886** | 154 | ERF | AT4G25480<br>(DREB1A) | .6HG0500460 |  | Regulator of responses to low<br>temperature and ABA signalling |
| **MLOC_61071** | 449 | Dof | AT3G52440 | .3HG0250060 |  | TF involved in regulation of<br>gene expression |
| **MLOC_54606** | 258 | WRKY | AT1G29860<br>(WRKY71) | 3HG0230420 |  | Regulator of shoot branching<br>and flowering time |
| **<u>MLOC_63184</u>** | 220 | WRKY | AT4G31550<br>(WRKY11) | 2HG0130750 |  | Regulator of ABA-dependent<br>stress responses |
| **<u>MLOC_10556</u>** | 138 | MYB | AT4G32730<br>(MYB3R-1) | 3HG0206530 |  | MYB TF involved in<br>transcriptional regulation |
| **MLOC_75795** | 187 | NAC | AT4G29230<br>(SND4) | 3HG0196360 |  | Negative regulator of flowering<br>and secondary cell wall<br>biosynthesis |
| **MLOC_5375** | 613 | MIKC_MAD<br>S | AT4G18960<br>(AG) | 3HG0202320 | -3,10 | MADS-box TF involved in the<br>development of floral organs |
| **MLOC_57142** | 834 | GATA | AT1G51600<br>(ZML2) | 4HG0283820 | -1,01 | Stress-inducible TF involved in<br>ABA-, drought- and salt-<br>response |
| **<u>MLOC_37446</u>** | 86 | G2-like | AT3G46640<br>(PCL1) | 3HG0273500 | -2,09 | Regulator of circadian clock<br>oscillation |

Underlined TFs were also identified as WT-specific in comparisons with all four SL mutants. \*among WT-specific drought-induced DEGs (3,811)

### Concluding model: SLs as integrators of hormonal sensitivity, transcriptional control, and physiological drought acclimation

Taken together, the physiological, hormonal, and transcriptomic datasets support a coherent model in which SLs mainly regulate drought response quality rather than response magnitude (Fig. 7). Across different mutants and genetic backgrounds, disruption of SL biosynthesis or perception consistently led to impaired maintenance of leaf water status, increased PSII photoinhibition, and transcriptional hyperactivation, indicating a higher stress burden. Despite strong activation of ABA metabolism in all genotypes, SL mutants showed signs of reduced ABA responsiveness, including changes in stomatal behaviour, weakened activation of ABA-dependent transcription factors, and inadequate induction of protective metabolic pathways. At the transcriptomic level, this resulted in repression of photosynthesis and carbon metabolism, along with activation of DNA repair and cell cycle processes, reflecting a shift from adaptive acclimation to damage-related responses. The repeated appearance of ABA-related TF modules across various datasets, including ERF/DREB, WRKY, NAC, and MYB regulators, further suggests that SL signalling allows the selective activation of regulatory nodes that control ROS homeostasis, wax deposition, and structural reinforcement. Overall, these findings support a model where SLs boost drought tolerance by strengthening ABA-dependent transcriptional networks and maintaining physiological stability, thereby aligning developmental structure with stress-responsive gene regulation (Fig. 7).

**Figure 7.**
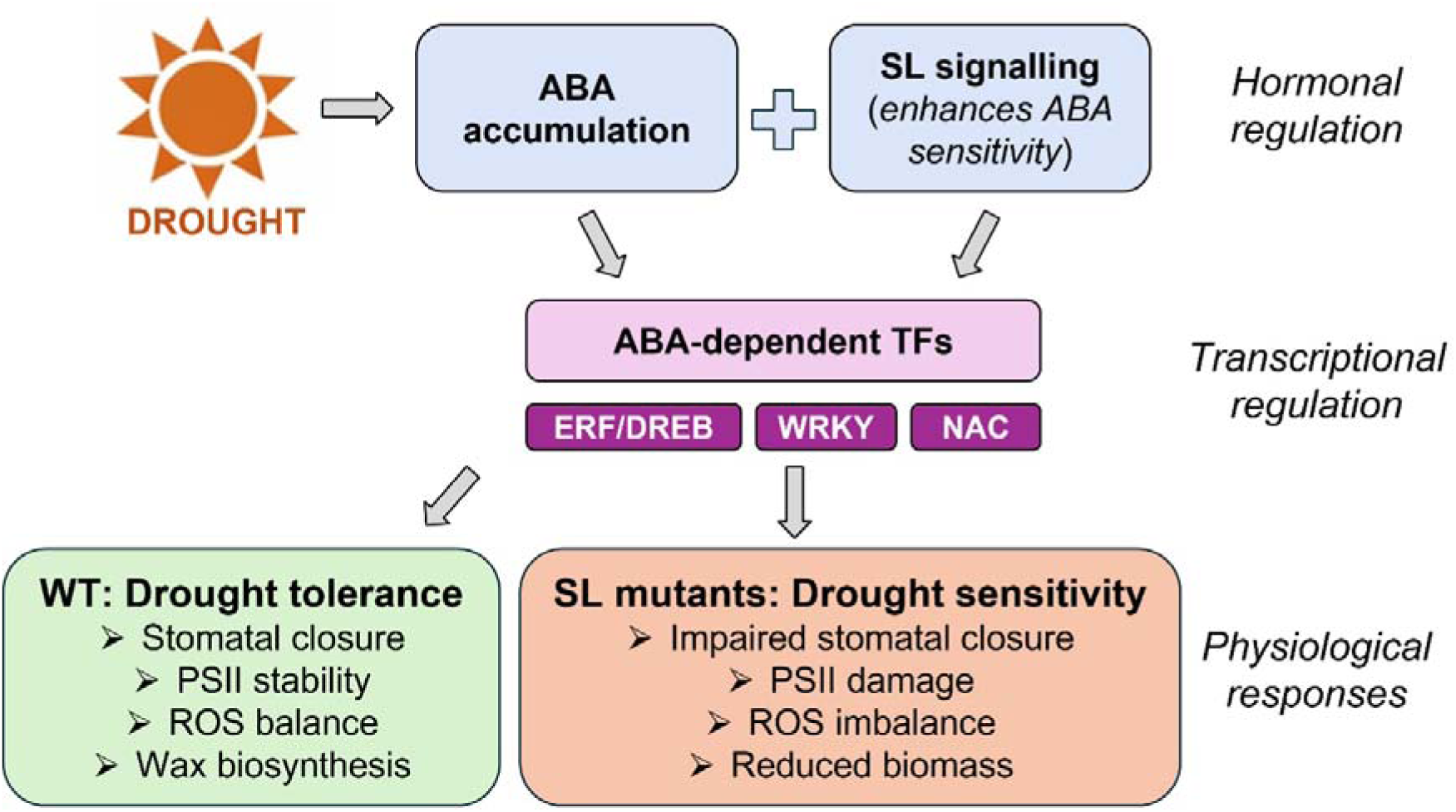
Proposed model of SL-mediated modulation of drought responses in barley. Schematic representation of the integrative role of SLs in shaping drought responses at hormonal, transcriptional, and physiological levels. Drought stress causes ABA accumulation in all genotypes; however, SL signalling enhances ABA sensitivity and promotes the activation of ABA-dependent TF modules, including ERF/DREB, WRKY, NAC, and MYB families. In WT plants, coordinated TF activation supports adaptive responses, including stomatal regulation, wax deposition, ROS homeostasis, and maintenance of PSII functionality, thereby improving drought acclimation. Conversely, SL mutants show reduced activation of ABA-dependent TF networks, along with transcriptional hyperactivation characterised by the repression of photosynthesis and metabolic pathways. This regulatory imbalance results in impaired stomatal control, oxidative stress, photoinhibition, and decreased biomass accumulation, collectively increasing drought sensitivity.

## Materials and methods

### Plant material, growth conditions and drought stress

For targeted mutagenesis, plant growth conditions were adapted from previous protocols (Hensel *et al*., 2009; Marthe *et al*., 2015), with slight modifications. Barley (*Hordeum vulgare* L. cv. Golden Promise) grains were sown in trays filled with a substrate mixture consisting of compost soil (*Komposterde*), white and black peat (Klasmann Substrate 2), and sand in a 2:2:1 ratio. Germination occurred in a phytochamber set to 14/12□°C (day/night), with a 12-hour photoperiod and a photon flux density of 136□μmol□m□²□s□¹. Approximately two weeks after germination, seedlings were transplanted into 2□L pots (Ø 16□cm) and transferred to a controlled greenhouse maintained at 18/16□°C (day/night), with a 16-hour photoperiod and a photon flux density of 170□μmol□m□²□s□¹. At the start of the tillering stage (Zadoks/BBCH stage 29), each pot was supplemented with 15□g of Osmocote® Pro 3-4M (ICL Deutschland), a slow-release fertiliser. Donor plants used for *Agrobacterium*-mediated transformation were grown under these conditions until 12-14 days after pollination, when immature caryopses were harvested for embryo isolation. Primary regenerants obtained through transformation were transplanted into soil after rooting and cultivated under the same greenhouse conditions. WT Golden Promise plants and Cas9 endonuclease-derived mutant lines were cultivated until maturity for phenotypic and molecular analyses and seed production.

For the analysis of shoot architecture, a single seed was sown in a pot (13 × 13 × 13 cm) filled with garden soil. The analyses were carried out weekly throughout the plant’s growth period. All plants were grown in a growth chamber under controlled conditions (16/8 h photoperiod, 20°C, 420 μE m□² s□¹ light intensity).

To evaluate the effect of SL application on barley tillering, GR24^5DS^ (StrigoLab, Turin, Italy) was utilised. Four seeds per genotype were sown in pots (7.5 × 7.5 × 10 cm) filled with garden soil. Plants were irrigated daily with 50 ml of solution from 1 to 10 DAS and 100 ml from 11 to 18 DAS, containing either 10 µM GR24^5DS^ or 0.01% acetone as a control. A paired Student’s t-test was conducted to identify statistically significant differences between treatments. For each condition, a total of 16 plants (grown in four pots) were assessed.

The survival rate for all analysed genotypes was determined using the method described above, with minor modifications. In each pot, 15 seedlings per genotype were arranged in two rows (one for the tested mutant and one for the wild type). Watering was stopped at 10 DAS and withheld for 15 days, then resumed for a further 3 days. Survival was evaluated at 28 DAS based on a total of 60 plants per genotype grown across four pots. Statistical significance between genotypes was assessed using a paired Student’s t-test.

Drought stress was applied following the protocol described previously (Daszkowska-Golec *et al*., 2019), with minor modifications. In brief, 15 seeds were sown in boxes (400 mm × 140 mm × 175 mm) filled with a sandy loam and sand mixture. Eight boxes were prepared in total. Plants were cultivated in a greenhouse under controlled conditions (20/18°C day/night, 16/8 h photoperiod, 420 μE m□² s□¹ light intensity) and well-watered conditions (14% vwc) for the first 10 days after sowing (DAS). Afterwards, irrigation was withheld from four boxes per genotype to induce drought stress. By 15 DAS, when soil moisture dropped to 3%, the plants were transferred to a growth chamber set at 25/20°C (day/night), with a 16/8 h photoperiod and a light intensity of 420 μE m□² s□¹. Severe drought (3-1.5% vwc) was maintained for 10 days (16-25 DAS). Control plants were grown under identical conditions but with optimal watering (14% vwc). Soil moisture was monitored daily using a time-domain reflectometer (TDR) EasyTest (Institute of Agrophysics, Polish Academy of Sciences).

### Targeted mutagenesis

After sequencing the target genes in cv. Golden Promise, the online platforms DESKGEN (now discontinued), Benchling, and WU-CRISPR were used to identify potential target motifs (TMs) within their coding sequences. The final selection of TMs integrated platform-derived efficiency scores with additional criteria, such as the predicted 2D structure of the guide RNAs and off-target filtering via BLAST against the barley genome (MorexV3_pseudomolecules_assembly). Guide RNA expression cassettes were assembled using the CasCADE modular vector system (Hoffie et al., 2026). Complementary oligonucleotides corresponding to the target motifs were annealed and cloned into BsaI-digested entry vectors carrying the *Triticum aestivum U6* (*TaU6*) promoter. Depending on the target gene, two (*HvD10*) or four (*HvD17*, *HvMAX1A*; *HvD14*) gRNA modules were assembled into a single intermediate vector via Esp3I-mediated Golden Gate cloning. This intermediate construct was then combined with the Cas9 expression unit and an auxiliary module using BsaI. Finally, all expression units were transferred into a binary vector - containing a hygromycin resistance gene (*hpt*) for plant selection - by SfiI digestion and ligation. All plasmid constructs were propagated in *E. coli* and transferred into *A. tumefaciens* strain AGL1 by electroporation, prior to DNA transfer to immature embryos of spring barley cv. Golden Promise, as described previously (Hensel *et al*., 2009). Briefly, embryos were excised from caryopses 12-14 days after pollination and co-cultivated for 48 hours with the *A. tumefaciens* strain AGL1 carrying the respective binary vector. Explants were then transferred to a selective medium containing Timentin and hygromycin for callus induction and plant regeneration. After DNA extraction from primary regenerants, mutation screening was performed by PCR amplification of target regions, followed by Sanger sequencing and sequence analysis. Transgene presence was assessed by PCR using primers specific for *cas9* and *hpt* (Supplementary Table S12). The plants carrying mutations were advanced through segregating generations and repeatedly genotyped to identify individuals that were homozygous for the desired mutation(s) and free of the transgenes. When possible, transgene-free and homozygous mutant lines were used for downstream analyses.

#### Transcriptomic analysis

RNA was extracted from four biological replicates, each consisting of 2 cm segments from the second leaf, taken 3 cm below the tip and pooled from three separate plants, using the mirVana miRNA Isolation Kit (Thermo Fisher Scientific, catalogue number AM1560). Library preparation and sequencing (150-nt paired-end reads) were conducted on the Illumina NovaSeq 6000 platform by Novogene Genomics Service (Cambridge, UK), which also processed the initial data using their RNA-seq analysis pipeline. Genes with an adjusted p-value < 0.05 and a log2 fold change ≥ 1 or ≤ -1 were classified as differentially expressed.

#### Gene Ontology enrichment analysis

Gene Ontology (GO) enrichment analysis was performed using ShinyGO (version 0.85.1). Analyses were carried out using the GO Biological Process database. An adjusted false discovery rate (FDR) threshold of 0.05 was applied to identify statistically significant enrichment. For each analysis, the top 20 significantly enriched GO terms were selected for further interpretation. Default background gene sets provided by ShinyGO were used unless otherwise stated.

#### Promoter sequences analysis and identification of TF

To analyse promoter sequences, 1500 bp upstream of the start codon (“Flank Gene”) of the differentially expressed genes (DEGs) was retrieved from the *Hordeum vulgare* TRITEX gene (Morex_V2_scaf) dataset using the BioMart tool (https://plants.ensembl.org/index.html). These sequences were then used as input for the ‘Regulatory prediction’ tool available at PlantRegMap (https://plantregmap.gao-lab.org/) to identify potential transcription factor (TF) binding sites and regulatory relationships. The analysis also included identifying TFs with significantly enriched targets within the DEG set. Arabidopsis homologues of the detected barley TFs were determined using the Plant Transcription Factor Database (https://planttfdb.gao-lab.org/).

#### Phytohormone analysis

Material for phytohormone analysis was prepared from young leaves following lyophilisation. Approximately 10 mg of lyophilised leaf tissue was placed into 2 ml safe-lock tubes (Eppendorf, Germany). Empty tubes served as blanks. Before extraction, two 3 mm ceria-stabilised zirconium oxide beads were added to each tube. Extraction and purification were performed as previously described (Šimura *et al*., 2018), with minor modifications. Briefly, 1 ml of ice-cold 50% aqueous (v/v) ACN containing internal standards (final concentration 100 nM) was added to each tube. Samples were homogenised for 5 min at 27 Hz in a mixer mill (MM 301, Retsch, Germany), then sonicated for 3 min at 4 °C in an ultrasonic bath (Sonorex, BANDELIN, Germany). Extraction continued for at least 30 min on an overhead shaker (Reax 32, Heidolph, Germany), followed by centrifugation for 10 min at 14,000 rpm and 4 °C (CT 15 RE, Himac, Japan). The resulting supernatant was transferred to new Eppendorf tubes (Germany) and purified using polymer-based SPE cartridges (Oasis PRIME HLB, Waters, USA; 1 cc per 30 mg). After loading, the flow-through was collected, and the remaining phytohormones were eluted with 1 ml of 30% (v/v) ACN; both fractions were combined. Lastly, samples were dried at 40 °C in a vacuum concentrator (RVC 2-33 IR, Martin Christ, Germany) and stored at -20 °C until analysis.

Absolute quantification of all targeted phytohormones, except gibberellins, was carried out using an Acquity Premier Binary Solvent System coupled to a Xevo TQ-S cronos mass spectrometer with UniSpray/ZSpray™ ionisation (UPLC-ESI-MS/MS; Waters, USA). Gibberellins were analysed on a Vanquish UHPLC system coupled to a Q Exactive Plus mass spectrometer with a heated electrospray ionisation source (UHPLC-HESI-HRMS; Thermo Scientific, USA). Spectra were acquired either in MRM mode (Xevo) or FullMS/dd-MS² mode (Orbitrap). Data were processed with MassLynx v4.2 SCN 1048 (Waters, USA) or TraceFinder v4.1 (Thermo Scientific, USA). Calibration curves (12 points, 0.5-1000 nM) were prepared from standard mixtures, using normalized peak areas of □M+H]□/□M-H]□ ions extracted from chromatograms. Linear regression (least squares) was applied to evaluate curve linearity. Compound identification in plant extracts was based on retention time, exact m/z, and isotope pattern, compared with authentic standards. MS² spectra were also matched against a custom spectral library for confirmation. Absolute quantification was performed as previously described (Eggert and Von Wirén, 2017; Milyaev *et al*., 2022).

Methanol (MeOH) and acetonitrile (ACN) were purchased from Th. Geyer GmbH & Co. KG (Germany), while formic acid (FA) was obtained from Biosolve Chimie (France). All mixtures were prepared using deionized distilled water from the Milli-Q® Reference System (Merck, Germany), which also served as the mobile phase for chromatography. Phytohormone standards, along with isotopically labelled internal standards, were purchased from OlChemim s.r.o. (Czech Republic) or Merck KGaA (Darmstadt, Germany).

## CRediT authorship contribution statement

Conceptualization: MMa; Methodology: JK, MMe, GH, ADG, MMa; Investigation: IMF, WB, MMe, ADG, MMa; Writing - original draft: IMF, WB, JK, MMe, GH, ADG, MMa; Writing - review & editing: IMF, WB, JK, MMe, GH, ADG, MMa; Funding acquisition: MMe, MMa. All authors read and approved the manuscript.

## Ethics approval and consent to participate

Not applicable.

## Consent for publication

Not applicable.

## Declaration of Generative AI and AI-assisted technologies in the writing process

During the preparation of this work, the authors did not use AI tools.

## Acknowledgements

This study was supported by the National Science Centre, Poland (2018/31/F/NZ2/03848) and German Research Foundation (ME 3356/7-1). We thank Sabine Sommerfeld for technical assistance during transformation and Dr. Yudelsy Antonia Tandron Moya for performing phytohormone measurement.

## Availability of data and materials

Transcriptomic data: ArrayExpress repository E-MTAB-17039.

Reviewer link: https://www.ebi.ac.uk/biostudies/arrayexpress/studies/E-MTAB-17039?key=97af5377-3716-415d-97b1-6c34efbc646e

Plant material will be made available on request.

## Supplementary Information

**Table S1. Summary of mutations detected in the generated mutant lines**. Insertions and deletions (indels) are indicated in base pairs (bp) relative to the reference sequence. Although individual indels occasionally correspond to multiples of three nucleotides, the combined mutations within each allele resulted in frameshift mutations leading to predicted truncated proteins.

**Table S2**. DEGs identified by RNA-seq in individual SL mutants compared with the WT.

**Table S3**. DEGs were shared between high-tillering SL mutants and across all analysed SL mutants.

**Table S4**. Gene Ontology Biological Process enrichment analysis of shared DEGs in SL mutants.

**Table S5**. TF enrichment and regulatory interactions associated with shared DEGs in SL mutants.

**Table S6**. Phytohormone profiling of WT and SL mutants under control conditions.

**Table S7.** DEGs identified by RNA-seq in individual SL mutants in response to drought stress.

**Table S8.** Overlap of drought-responsive DEGs among SL mutants and WT.

**Table S9.** Genotype-specific and shared drought-responsive DEGs in WT and SL mutants.

**Table S10.** Gene Ontology Biological Process enrichment of genotype-specific drought-responsive DEGs in WT and SL mutants.

**Table S11.** TF enrichment and predicted regulatory interactions associated with WT- and mutant-specific drought-responsive DEGs.

**Table S12**. Phytohormone profiling of WT and SL mutants under drought conditions.

**Table S13**. Primer sequences used in this study.

**Figure S1. Schematic representation of the target sites selected for Cas endonuclease-mediated mutagenesis in barley strigolactone-related genes.** Exon-intron structures of *HvD17*, *HvD10*, *HvMAX1a*, and *HvD14* are shown with the positions of sgRNAs used for targeted mutagenesis indicated by red dashed lines. Boxes represent exons, and black connecting lines indicate introns. Scale bar = 100 bp.

**Figure S2. Grain size and thousand-grain weight in barley SL mutants.** (A) Thousand-grain weight (MTZ), (B) grain length, and (C) grain width measured in WT and SL mutants. Data are presented as mean values with standard error (SE) (n=6 per genotype). Different letters indicate statistically significant differences between genotypes according to one-way ANOVA followed by Tukey’s HSD test (p < 0.05). (D-H) Representative images of grains from WT, *Hvd17-1, Hvd10-1, Hvmax1a-1*, and *Hvd14-1* illustrating differences in grain size and shape. Scale bars=1 cm.

**Figure S3. Specific changes in phytohormone metabolism in barley SL mutants.** (A) Abscisic acid (ABA), (B) Dihydrophaseic acid (DHPA), (C) phaseic acid (PA), and (D) trans-Zeatin (tZ) levels. Data are shown as mean values with standard error (SE) (n=4). Asterisks denote significant differences between mutants and the wild-type (WT) according to Student’s t-test (*p*-values indicated by \**p*<0.05, \*\**p*<0.001). Phytohormone concentrations are expressed in ng gLJ¹ of dry weight (DW).

**Figure S4. Drought-induced changes in PSII performance in barley SL mutants.** (A) Maximum quantum efficiency of PSII (φPLJ, equivalent to Fv/Fm). (B) Performance index on absorption basis (PI ABS), reflecting overall PSII functionality. (C) Absorption flux per active reaction centre (ABS/RC), indicating changes in the functional status of PSII reaction centres. (D) Dissipated energy flux per cross section (DILJ/CSLJ), reflecting enhanced non-photochemical quenching under drought stress. Data are presented as box plots showing median, interquartile range, and minimum/maximum values; individual data points represent measurements from single plants (n = 16 per genotype and treatment).

