## Supplementary Figures for "Generation and characterization of a barley strigolactone mutant collection: from plant architecture to drought stress response"

### Slide 1
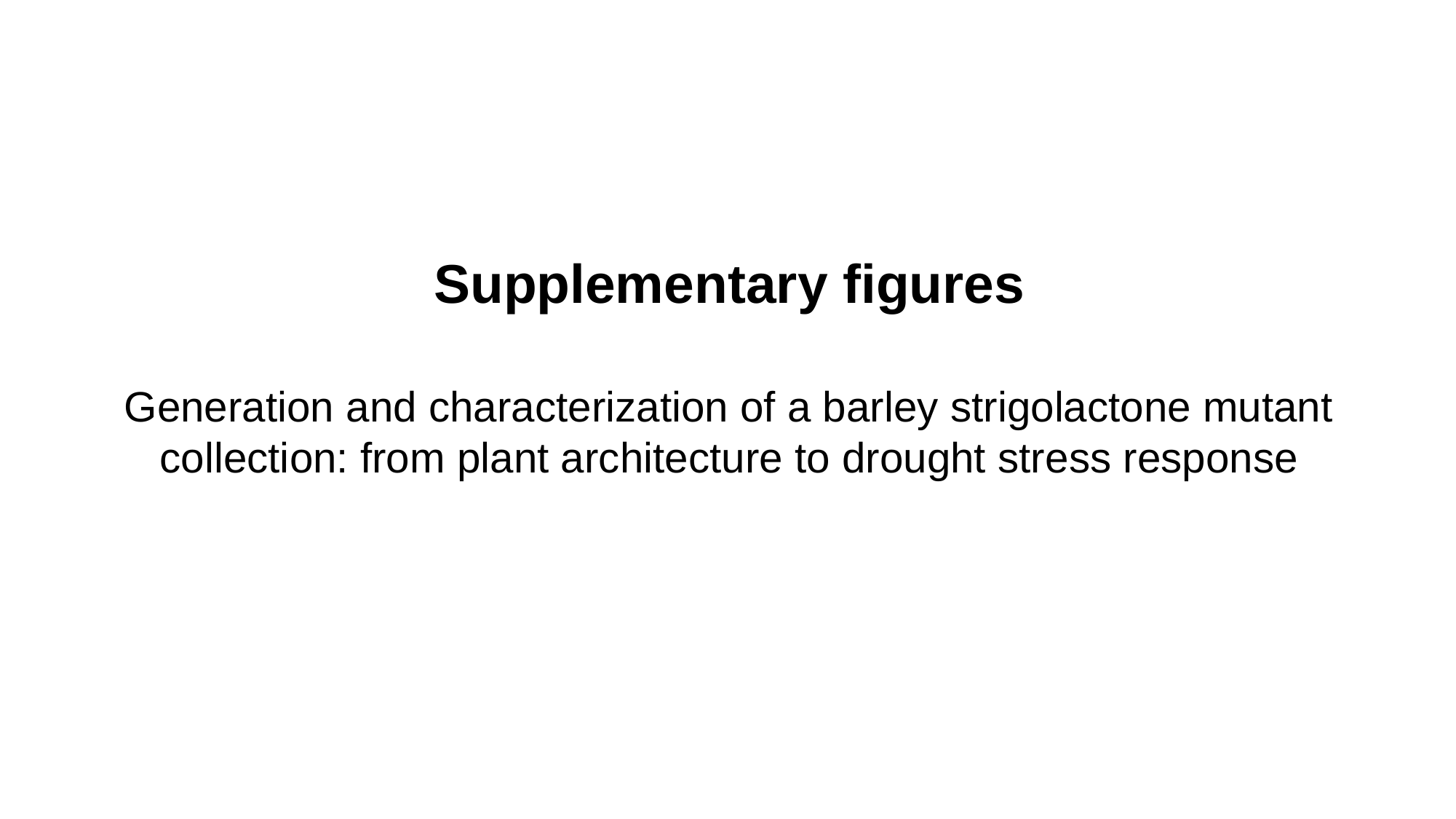

Supplementary figures
Generation and characterization of a barley strigolactone mutant collection: from plant architecture to drought stress response

### Slide 2
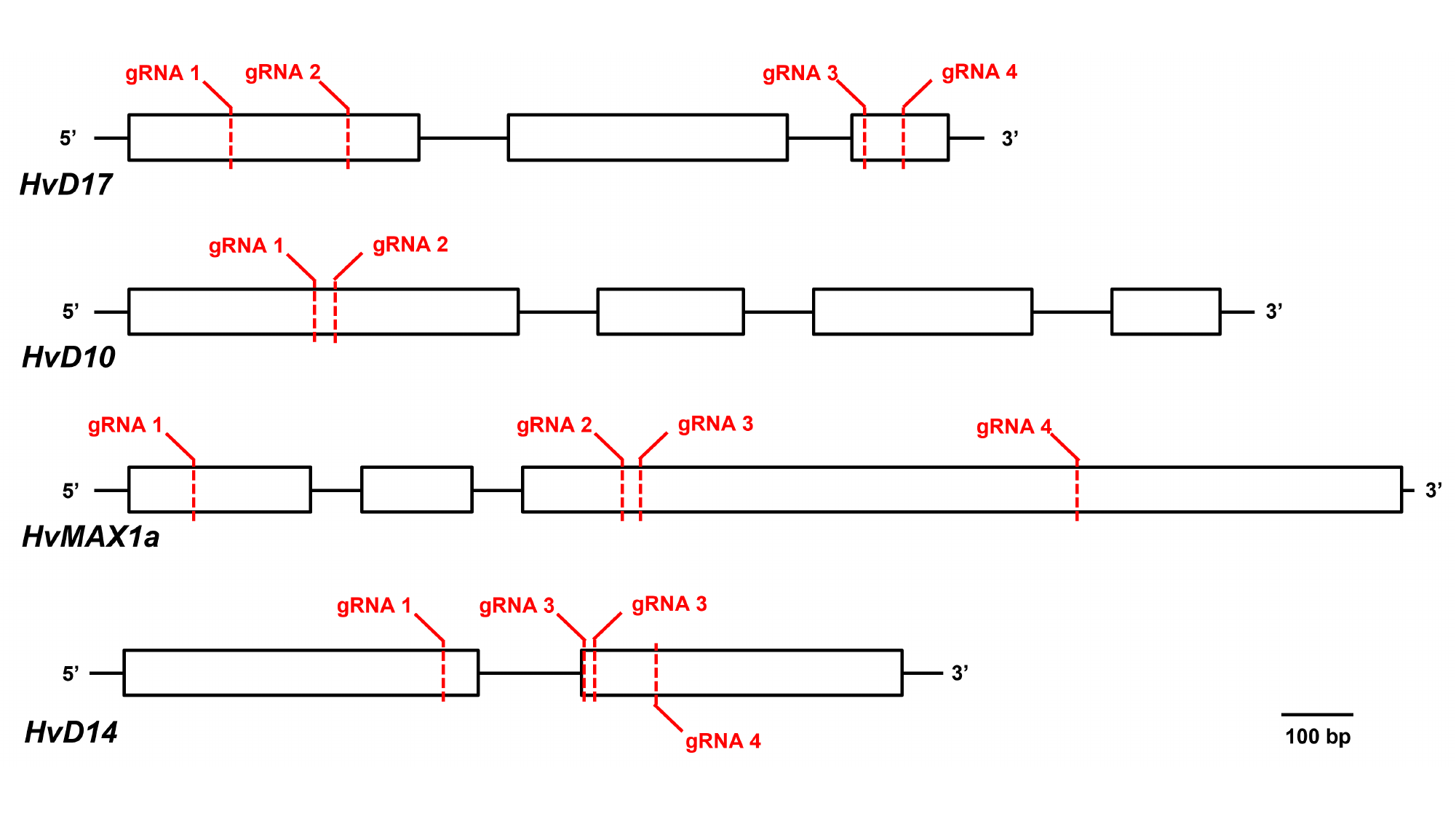

### Slide 3
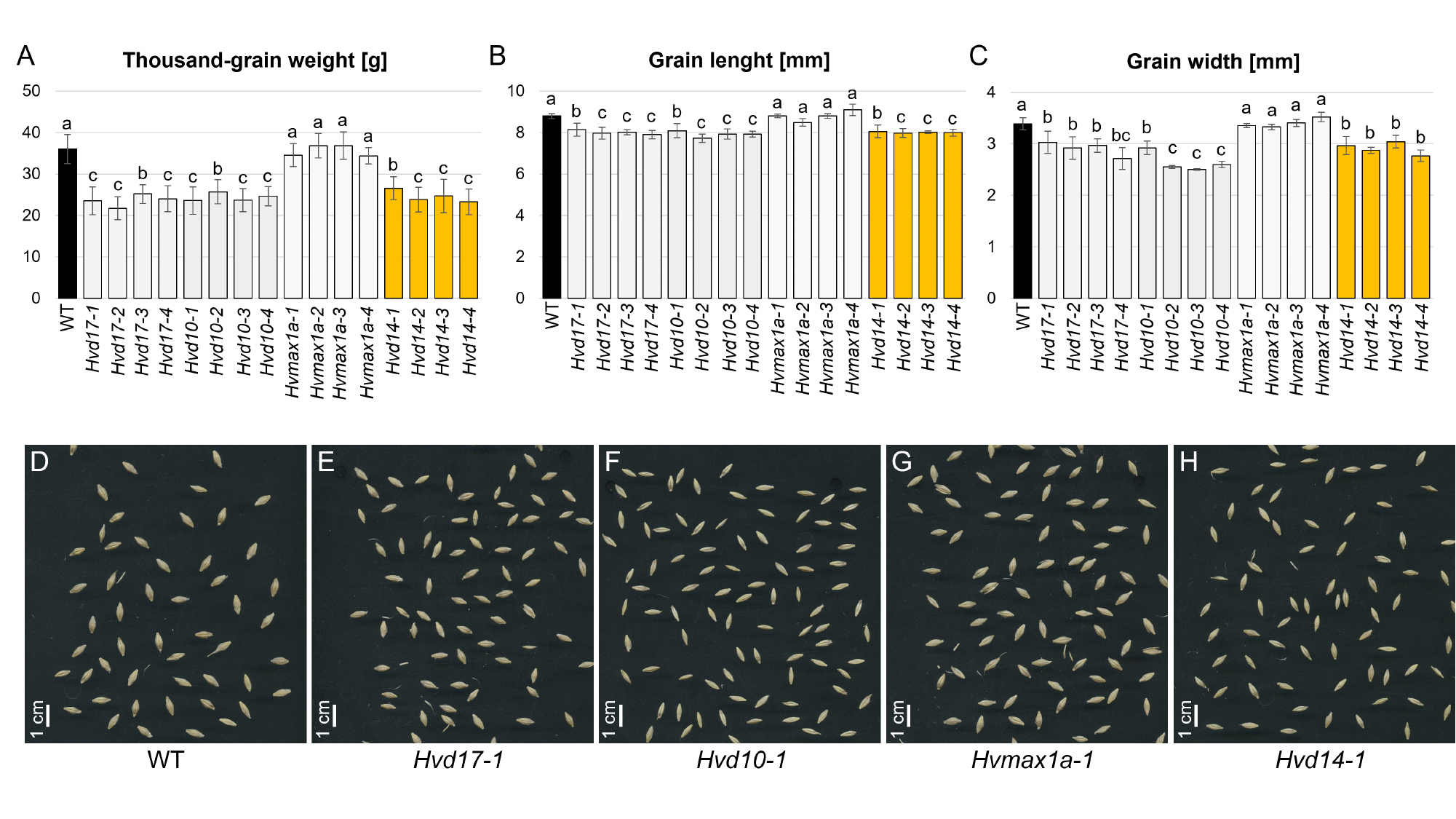

### Slide 4
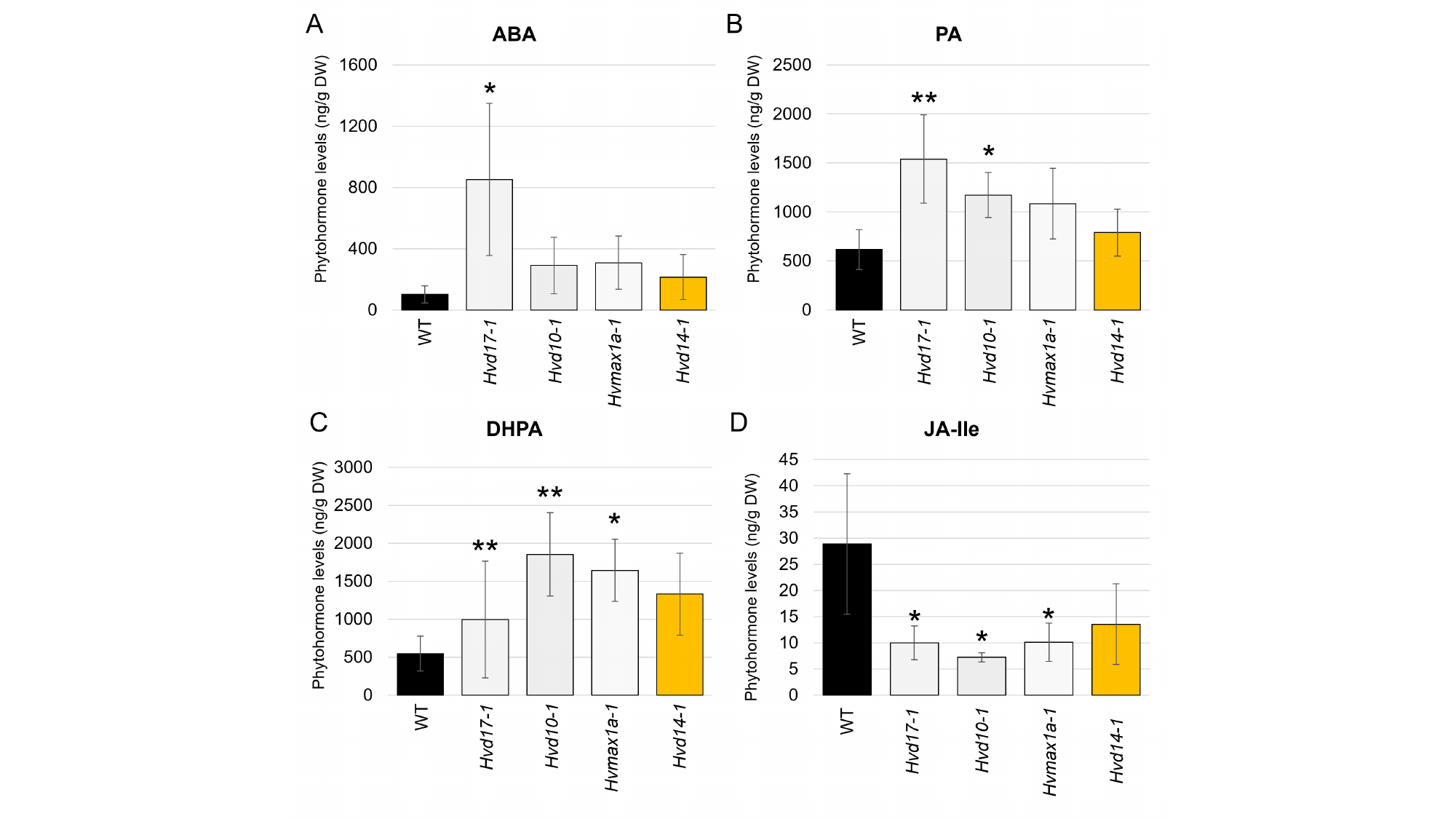

### Slide 5
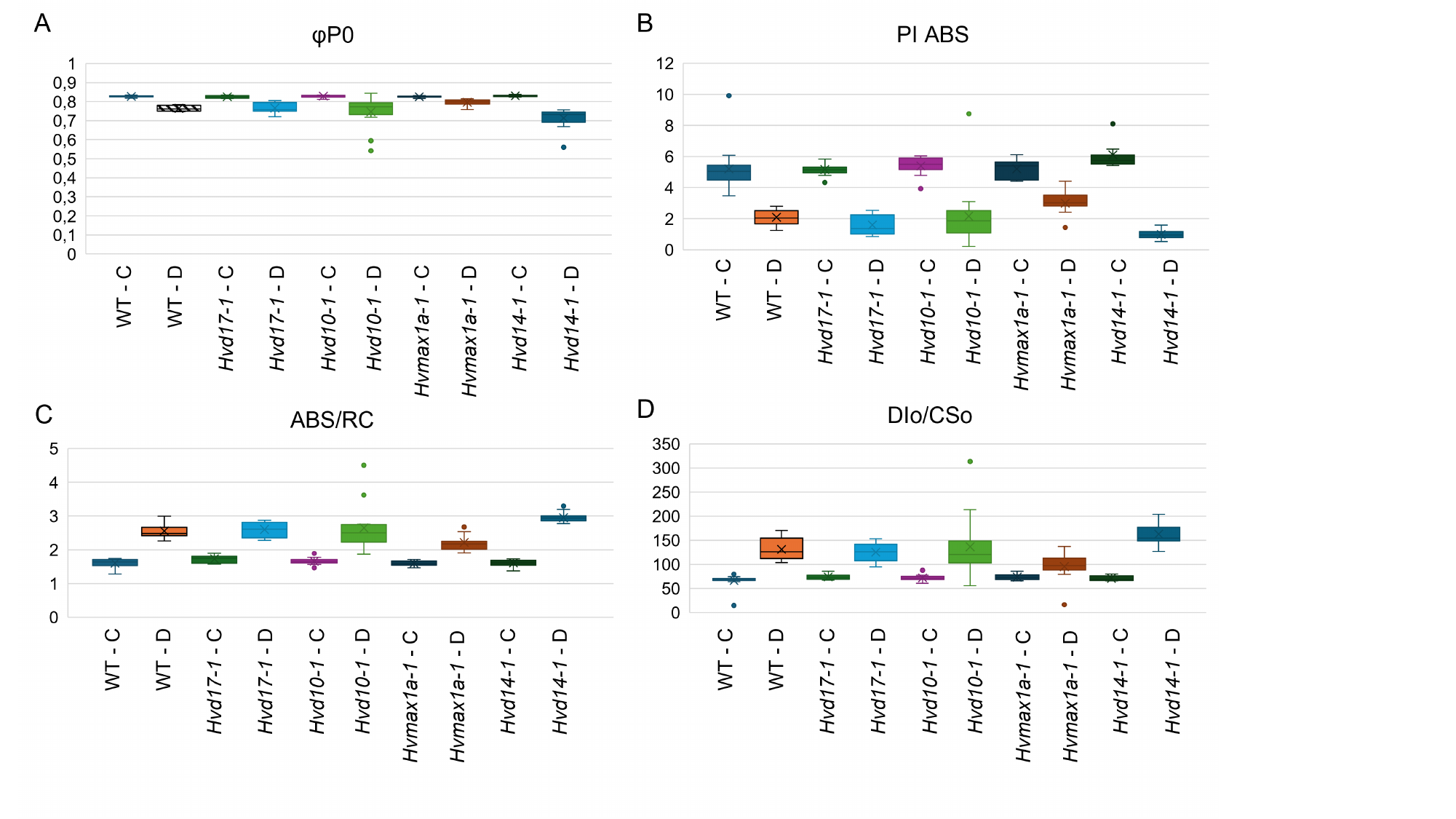
